# Evolution of the RNA alternative decay *cis* element into a high-affinity target for the immunomodulatory protein Roquin

**DOI:** 10.1101/2024.07.30.605734

**Authors:** Jan-Niklas Tants, Katharina Friedrich, Jasmina Neumann, Andreas Schlundt

## Abstract

RNA *cis* elements play pivotal roles in regulatory processes, e.g. in transcriptional and translational regulation. Two stem-looped *cis* elements, the constitutive and alternative decay elements (CDE and ADE, respectively) are shape-specifically recognized in mRNA 3’ untranslated regions (UTRs) by the immune-regulatory protein Roquin. Roquin initiates mRNA decay and contributes to balanced transcript levels required for immune homeostasis. While the interaction of Roquin with several CDEs is described, our knowledge about ADE complex formation is limited to the mRNA of *Ox40*, a gene encoding for a T-cell co-receptor. The *Ox40* 3’UTR comprises both a CDE and ADE, each sufficient for Roquin-mediated control. Opposed to highly conserved and abundant CDE structures, ADEs are rarer, but predicted to exhibit a greater structural heterogeneity. This raises the question how and when two structurally distinct *cis* elements evolved as equal target motifs for Roquin. Using an interdisciplinary approach we here monitor the evolution of sequence and structure features of the *Ox40* ADE across species. We designed RNA variants to probe en-detail determinants steering Roquin-RNA complex formation. Specifically, those reveal the contribution of a second RNA-binding interface of Roquin for recognition of the ADE basal stem region. In sum, our study sheds light on how the conserved Roquin protein selected ADE-specific structural features to evolve a second high-affinity mRNA target *cis*-element relevant for adaptive immune regulation. As our findings also allow expanding the RNA target spectrum of Roquin, the approach can serve a paradigm for understanding RNA-protein specificity through back-tracing the evolution of the RNA element.

## INTRODUCTION

RNA elements regulate numerous cellular processes, such as gene silencing, translation, mRNA decay and localization, or integrate external stimuli. While the first discovered so-called *cis*-regulatory elements were single-stranded, e.g. mRNA start codons^1^ and AU-rich elements (ARE)^2, 3^, we continue identifying folded elements comprising a great structural variety^4–6^. Hairpin elements account for a large number of these structures, and they occur in different flavours with varying loop sizes or internal bulges and branches. Tertiary and long-range contacts in RNAs are rare^7^, but crucial as they mediate formation of highly complex structures, e.g. in viral replication elements^8^. Evolutionarily, these critical contacts are often highly conserved^9^. RNA *cis* elements serve as a binding platform for *trans*-acting factors, e.g. proteins that confer functionality to the protein RNA complex (RNP). RNA target recognition can be either sequence-^10^ or shape-specific^11^. Several proteins exploit their modular design to integrate both sequence and shape specificity^12, 13^, or to increase affinity towards their targets^14^ via multiple RNA-binding domains. The limited number of tertiary RNA contacts favours formation of isolated structured building blocks that often form *cis* element hubs and can exert cooperative or antagonistic effects^15, 16^. Changes in *cis* element sequence or structure have been associated with diseases, as subtle alterations can lead e.g. to disruption of functional RNPs^17^.

RNA shape recognition requires a precise/defined geometry, as sequence-specific contacts between protein and RNA are limited because RNA structure is accompanied by shielding nucleobases^18^. Complex formation is then depending on recognition of exposed single-stranded nucleotides as found in loops or kinks^18^, or the overall RNA shape. By now there are only a few RNPs that provide specific recognition of *cis* element shape. Most prominent are U1A^19^, FUS^12^, double-stranded RNA-binding proteins (dsRBPs)^20, 21^ and Roquin^22^. Roquin controls transcript levels through initiation of mRNA decay^23^. It engages with hairpin structures that comprise tri- and hexaloops, which were termed constitutive and alternative decay elements (CDE and ADE, respectively)^22, 24^. The Roquin coreROQ domain harbours the stem-loop binding interface, called A-site, which is highly conserved across vertebrates, while a second interface (B-site) composed of flanking helical extensions of the ROQ domain (extROQ) was suggested to engage with double-stranded RNA^25^. CDEs and ADEs occur solitary or in clusters in mRNA 3’ untranslated regions (UTR)^16, 26^. At least 13 functional CDEs have been confirmed in nine 3’UTRs^16, 23, 24, 27, 28^ while only four ADEs^16, 24, 27^ were discovered so far. Predictions of further ADEs, however, indicated a great structural variety in contrast to the seemingly more conserved CDE fold^28^. Currently, we do not understand how Roquin integrates this structural variety of targets into high-affinity RNPs.

Both decay elements have proven sufficient for Roquin-mediated posttranscriptional gene regulation, but they can also act cooperatively, e.g. in *NFKBID*^16^. For the *Ox40* (*TNFRSF4*) tandem ADE-CDE cassette it was shown that Roquin engages with both elements independently to form a 2:1 complex with low nanomolar affinities^26^. Evolution of two functionally redundant elements might serve as a backup strategy for tight regulation of critical genes. It could also offer yet undiscovered hidden contributions to fine-tuning of gene regulation, which might be coupled to the structural differences between ADE and CDE. Evolution of a tri- and hexaloop RNA hairpin element as equal high-affinity targets for a single protein highlights the great plasticity of Roquin RNPs. Protein-RNA pairs of high functional relevance often co-evolved, meaning that sequence and structure alterations in one binding partner entailed adaptations in the other interacting molecule^29, 30^. One driving force for such co-evolution can be the affinity between the involved interaction partners^31^. For RNA, its poor structural predictability hampers a reliable judgement of preservation or modulation of RNA element structure upon changes in sequence throughout evolution^32, 33^, and it requires sophisticated bioinformatic approaches to also take co-variation into account^34^. Co-evolution of *cis*-*trans* pairs has led to specialised binding interfaces^35^, implying beneficial functions for the organism, and was found to occur even over comparably short time periods^36^. In fact, host pathogen interactions have been identified as a hotspot for rapid co-evolution of protein RNA pairs^37^. Consequently, the immune system is expected to reflect evolutionary adaptations on the protein and RNA levels

Roquin controls many targets with immune-regulatory functions. Up to date we lack information when and how the two types of decay elements evolved. The CDE was described first, however, no study so far analyzed whether ADE and CDE evolved simultaneously or sequentially. In the latter case, this could point at additional functions of the second, structurally different element required in immune system control. Since comparative analyses of *cis* elements often focus on mammals or even apes, we do have no examples of progenitor ADEs. Early ancestors of known ADEs and CDEs could deepen our knowledge of Roquin-RNA interaction, contribute to Roquin target definition by revealing RNA features key for recognition and shed light on the benefits of the evolution of two RNA elements similar in function, yet different in structure. Monitoring the evolution of an RNA *cis* element should reflect key steps in structure optimization towards a high-affinity target for a cognate RBP. It is thus tempting to track back the occurrence of decay elements. So far high-resolution structural studies of Roquin RNPs focused on small stem-loop elements bound by the coreROQ domain^22, 24^, omitting the full network of interactions of the extROQ interface with extended RNA elements. The precise interplay of Roquin A and B-sites is thus unknown as well as their contribution to structure selection during evolution of target *cis* elements. Of note, there is evidence that both sites contribute to recognition of distinct RNA elements^25, 38^. For a full understanding of Roquin target specificity, both RNA binding interfaces and an extended *cis* element contexts need to be considered that exceeds apical stem-loops used so far. In fact, in this context the larger structural plasticity of ADEs compared to CDEs can be exploited to reveal structural features of Roquin RNP formation that may have been overseen so far.

We here assessed the degree of secondary structure in *Ox40* ADEs from six mammals as well as two putative ancestor ADEs from reptiles and birds. Mammalian ADEs showed a high conservation of stability and shape, which significantly enhanced Roquin binding compared to possible ancestral ADEs. Technical mutants of human and murine *Ox40* ADEs revealed that the Roquin B-site is particularly sensitive to sequence changes leading to a lowered stability of the central stem. Further, geometric hallmarks and asymmetric base distribution within the RNA stem fine-tuned affinity during evolution. We observed that the orientation and size of a central bulge affects RNP formation. Together, we give evidence for the divergent roles of and target requirements in the Roquin A and B-sites. Our evolutionary study highlights the formation of stable hairpin structures as a key event in the evolution of ADEs as Roquin targets, while technical mutants reveal geometric parameters for the fine-tuning of complex formation that may have developed during evolution. We suggest that the mammalian ADE has evolved as a second and equal Roquin target *cis* element for critical genes under the pressure of the immune system.

## MATERIAL AND METHODS

### RNA sequence alignment and ADE prediction

*Ox40* 3’UTR sequences were obtained from the UCSC Genome Browser^39^. For opossum, chicken and lizard predicted 3’UTR sequences were used. Tandem ADE-CDE alignment was performed with Clustal Omega^40^. Phylogenetic trees were obtained from alignments. For visualization of exon and 3’UTR conservation of genes the Multiz Alignment of 100 vertebrates from the UCSC Genome Browser was used. RNA secondary structure prediction was done with the RNAfold tool from the Vienna RNA Websuite^41, 42^. ΔG values were obtained from the same webserver. RNA constructs for *in vitro* transcription were designed to be similar in size and to contain the predicted central bulge as well as the lower stem adjacent to the bulge, whenever possible.

### Protein expression and purification

Murine protein constructs of coreROQ (aa 171-326) and extROQ (aa 89-404) were expressed and purified as described before^26^. Briefly, an LB pre-culture of *E. coli* BL21 (DE3) with 50 µg/mL Kanamycin was inoculated from a single colony and incubated at 37°C overnight shaking. From this, a preparative culture in M9 minimal medium containing 1 g/L ^15^N NH_4_Cl was started. Protein expression was induced with IPTG overnight, cells were harvested and lysed in 50 mM Tris pH 8.0, 500 mM NaCl, 4 mM β– mercaptoethanol. Proteins were purified via IMAC (Ni-NTA), subsequent TEV-cleavage and a reverse IMAC. After SEC (HiLoad S75 16/600 from GE) proteins were flash frozen, stored at −80°C and buffer exchanged to 150 mM NaCl, 20 mM Tris pH 7.0, 2 mM TCEP before usage. Purity was confirmed by SDS-PAGE and NMR spectroscopy.

### RNA *in vitro* transcription and purification

ADE RNAs (**Table 1**) were *in vitro* transcribed from a plasmid containing the HDV ribozyme for 3’-end homogeneity. After transcription by T7 polymerase at 37°C RNAs were purified via denaturing 12% polyacrylamide gels and eluted from gels by ‘crush-and-soak’. Purified RNAs were stored at −20°C in water. Quality was assessed by denaturing PAGE and NMR spectroscopy. For buffer exchange and concentrating Amicon® filter units were used. All RNAs were snap-cooled for folding prior to use, i.e. RNAs were heated to 95°C for 5 min and rapidly cooled down in an ice bath.

**Table 1:**
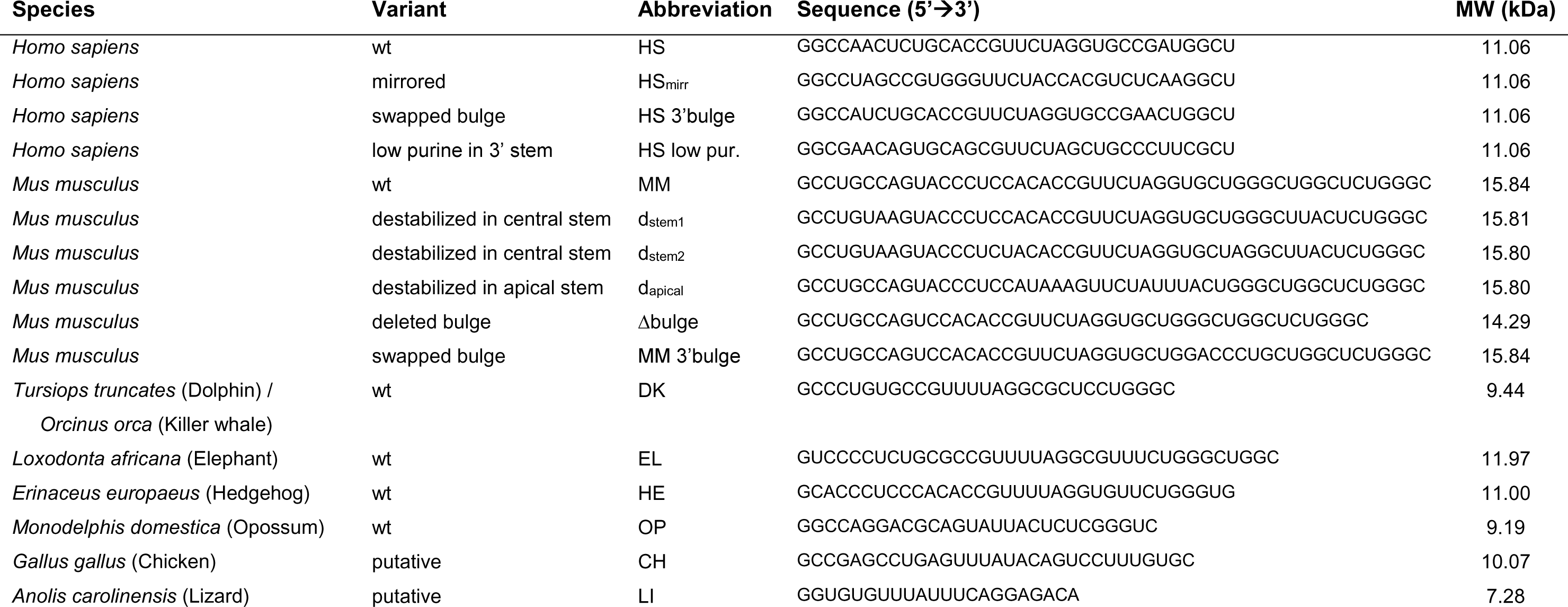
ADE RNAs used in this study. Corresponding abbreviations used are given. wt = wildtype; MW = molecular weight.

### NMR spectroscopy

All measurements were performed at Bruker AV spectrometers (600, 700, 900 MHz proton Larmor frequencies) equipped with triple-resonance cryoprobes. Spectra were processed with Topspin 4.0.6 and analyzed with NMRFAM-Sparky 1.414^43^. RNA only samples were buffer exchanged to 25 mM potassium phosphate pH 7.0, 50 mM potassium chloride while protein and RNP samples were dialyzed to 20 mM Tris pH 7.0, 150 mM NaCl, 2 mM TCEP prior measurement. For RNA imino proton assignments, ^1^H,^1^H-NOESY experiments were recorded at 283 K with mixing times of 150 or 200 msec. 1D imino proton spectra were recorded at 283, 298 and 310 K. Additionally, spectra in presence of 1 mM MgCl_2_ were recorded at 298 K. To monitor protein RNA interaction, ^1^H,^15^N-HSQC spectra were recorded of apo protein and a 1.5-fold excess of RNA at 298 K.

### Small-angle X-ray scattering (SAXS)

RNAs were dialyzed to 25 mM potassium phosphate pH 7.0, 50 mM potassium chloride with optional addition of 1 mM MgCl_2_, refolded by snap-cooling and flash frozen for transportation. Measurements were carried out at beamline P12 (PETRA III) at DESY, Hamburg, at 293 K. The ATSAS^44^ package and RAW^45^ 2.3.0 were used for data processing and analysis. R_g_s were obtained from RAW/ATSAS. Dimensionless Kratky plots were generated with RAW. RNA tertiary structure models were generated with RNAMasonry^46^ in 50 simulation steps based on our SAXS curves, and the fit quality to the experimental SAXS data was estimated with the implemented CRYSOL tool as Χ^2^.

### Circular dichroism (CD) spectroscopy

CD spectra were recorded in a JASCO J-810 spectrometer using 20 µM samples in 25 mM potassium phosphate pH 7.0, 50 mM potassium chloride in High Precision Cell Quartz cuvettes (Hellma Analytics) with a 1 mm path. Spectra were recorded in triplicates at 20 °C across wavelengths from 180 to 320 nm with a data interval of 0.5 nm. Melting curves were recorded from 5 to 95 °C at 260 nm in an interval of 0.2 °C with a temperature ramp of 1 °C/min. Melting curves were normalized to 5 °C. Data was plotted and melting points were calculated with OriginPro 2024 using a bi-dose response curve fit covering the full temperature range.

### Electrophoretic mobility shift assays (EMSA)

For gel shift assays RNAs were 5’-fluorescently labeled^26^. First, RNAs were dephosphorylated using Quick CIP (NEB) at 37 °C for 1.5 h. After phenol/chloroform extraction and ethanol precipitation, RNAs were phosphorylated using ATPγS (Cayman Chemical) and T4 PNK (NEB) at 37 °C overnight. Again, RNAs were phenol/chloroform extracted, precipitated and resuspended in 25 mM HEPES pH 7.4. RNAs were labeled at room temperature for 4 h in the dark by adding 5’-(iodoacetamido)fluorescein (IAF) (ThermoFisher) dissolved in DMSO to a final concentration of 7.5 mM. Labeled RNAs were purified via denaturing PAGE as described above and refolded in 20 mM Tris pH 7.0, 150 mM NaCl, 2 mM TCEP as before. For EMSAs, 2 µl RNA were incubated for 10 min at room temperature with 0.6 µg baker’s yeast tRNA_Phe_ (Roche), 1 mM MgCl_2_ and increasing amounts of protein in a volume of 20 µl. Protein concentrations of 0, 100, 200, 400, 700, 1,000, 2,000, 3,000, 5,000, 20,000 nM were used. Samples were mixed with loading buffer (1x TB, 60% glycerol, 0.02% Bromphenolbue) and run on 6% native polyacrylamide gels for 1 h as described before (^26^). Gels were imaged on a ChemiDoc Imager (Bio-Rad). For analysis ImageJ was used to quantify the free RNA. Fraction bound was calculated as before^26^. K_D_s are given as average plus standard deviation from individually fit triplicates using Hill1 fit in Origin.

## RESULTS

### Prediction of an *Ox40* ADE in multiple species

The *Ox40* 3’UTR provides posttranscriptional control over a critical T-cell co-receptor and is thus a key regulator within the mammalian immune system. Regulation occurs through Roquin binding to either or both ADE and CDE and we have previously shown that Roquin recognizes the two *cis* elements independently^26^. As typical for CDEs, the *Ox40* CDE shows a high degree of conservation among mammals (**Figure 1A**) with a near 100% sequence identity among apes and monkeys as assessed by a sequence alignment of the tandem ADE-CDE cassette. For all other species, only positions 1 and 2 within the loop showed variation as well as the 5’ nucleotide of the closing base pair. However, this position is always a pyrimidine and allows formation of a C-G or U-G base pair, conserving the overall hairpin fold with a triloop. All CDEs are predicted to adopt a six-base pair stem. In contrast, the ADE is prone to larger sequence variations, which is reflected in a larger score within the phylogenetic tree for the individual ADE compared to the CDE (**Supplementary Figure 1B** and **C**). As for the CDE, conservation within the ADE stem is high among apes and monkeys, but low for other species. Based on our sequence alignment, ape ADEs have an insertion in 5’ of the loop and show small deletions and sequence variations in 3’ of the loop compared to other mammals (**Figure 1A**). Interestingly, the ADE loop is more conserved than the CDE loop with only position four switching between C and U.

**Figure 1.**
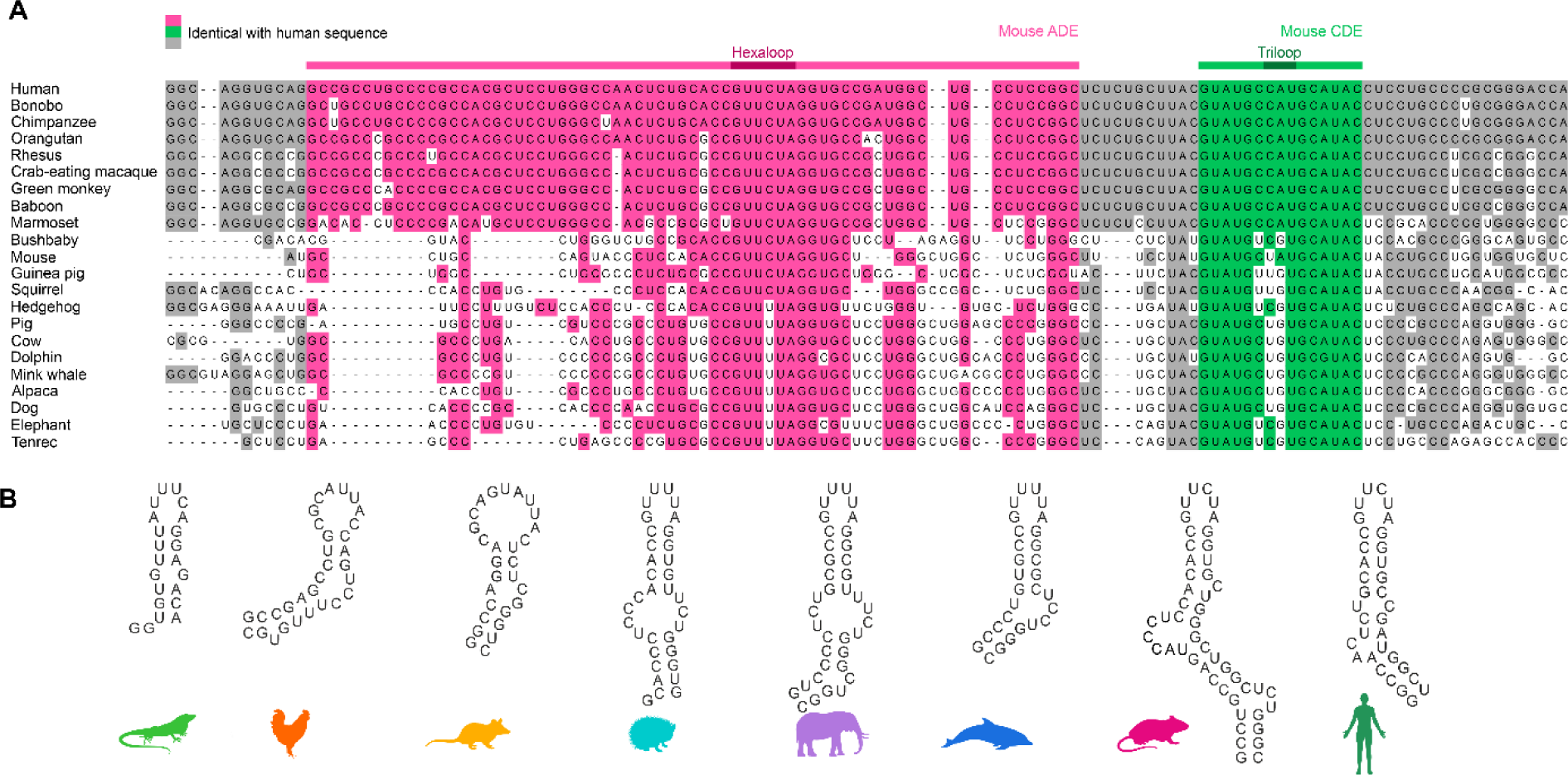
Evolutionary conservation of *Ox40* tandem ADE-CDE cassette. **A)** Sequence alignment of *Ox40* tandem ADE-CDE from multiple mammals. Bases identical with the human sequence are colored. Residues corresponding to the murine ADE and CDE are colored in magenta and green, respectively. Dark magenta and green highlight the hexa- and triloop, respectively. **B)** Secondary structure predictions of ADEs used in this study.

The high degree of CDE conservation suggests that it developed earlier, and that ADE evolution was coupled to advancements of the immune system. In line with this all species from the alignment showed a high sequence conservation for Roquin exons (**Supplementary Figure 2**), pointing at possible Roquin-CDE interactions throughout all mammals. This is supported by high sequence conservation of known Roquin target 3’UTRs, i.e. *NFKBIZ* and *NFKBID* (**Supplementary Figure 2**). Of note, chicken and lizard sequences aligned well with mammalian Roquin and target gene exons, while the corresponding 3’UTRs showed poor conservation. Assuming a similar immune-regulating function of Roquin in birds and reptiles, this could imply regulation of target genes by other *cis* element structures than the known CDE or ADE, like linear binding elements (LBEs)^16^ or AREs^47^. Roquin binding could also occur through conservation of a CDE/ADE fold despite high sequence variation. This could reveal possible progenitor ADEs and highlight structural features that were selected by Roquin during evolution. Predictions of mammalian ADEs (**Supplementary Figure 1B**) suggested formation of stable hairpin structures.

### *Ox40* ADEs adopt a stable fold

We designed ADE constructs of four mammals (**Table 1**, **Figure 1B**) based on the sequence alignment to assess critical structural ADE features. The high degree of conservation suggests (**Figure 1A** and **Supplementary Figure 1B**) that these are indeed functional ADEs in their respective species. We added an ADE-like sequence from an early mammal, i.e. the opossum, which showed poor conservation in the *Ox40* 3’UTR (**Supplementary Figure 2**). Additionally, we included analogous RNA regions as potential ADE ancestors in birds and reptiles to analyze their fold and stability. Therefore, we designed two putative ADEs from the aligned *Ox40* 3’UTR of chicken and lizard. Defined sequences were guided by the mammalian ADEs, i.e. expected to form hairpin structures and contain a U-rich loop of a size of 6-10 nucleotides. All ADE constructs were designed to be similar in size to avoid charge-driven bias in RNP interaction studies and to start with a G to facilitate *in vitro* transcription (**Table 1**). To assess the degree of secondary structure we recorded ^1^H NMR imino spectra of the RNAs at three temperatures (**Figure 2A** and **Supplementary Figure 3A**). The putative chicken and lizard ADEs showed only a few and broad imino proton signals, suggesting no stable and/or single conformation. Mammalian ADEs on the contrary showed dispersed peaks, indicative of a stable fold. Interestingly, peaks are more dispersed and uniform in linewidth for dolphin, mouse and human than for the hedgehog or elephant. This points at potential structural changes within the mammalian development that led to a further stabilization of the ADE fold. We confirmed the secondary structure of the human ADE by NMR spectroscopy (**Supplementary Figure 3B**). For all ADEs, only minor changes in the imino proton pattern were observed across the tested temperatures (**Supplementary Figure 3A**), suggesting formation of stable structures. To corroborate our observations, we measured CD melting curves (**Supplementary Figure 4**). Mammalian ADEs yielded melting temperatures of 69.3 – 72.3 °C (**Figure 2B** and **Supplementary Table 1**), in line with the predicted low ΔG values (**Supplementary Figure 1D**). For chicken and lizard, we observed melting temperatures of 39.0 and 63.5 °C, respectively. The lowered melting temperatures can be accounted for by an overall increased AU-content of the RNAs compared to the mammalian ADEs or increased loop-size (chicken) (**Supplementary Figure 1D**). The increased thermal stability of mammalian ADEs agrees well with our observation of secondary structure formation in NMR experiments. Despite sequence and structure variations the mammalian *Ox40* ADEs are comparably stable, suggesting the melting temperature to be optimal for gene regulation of *Ox40*, e.g. through an RNA fold-recognizing protein.

**Figure 2.**
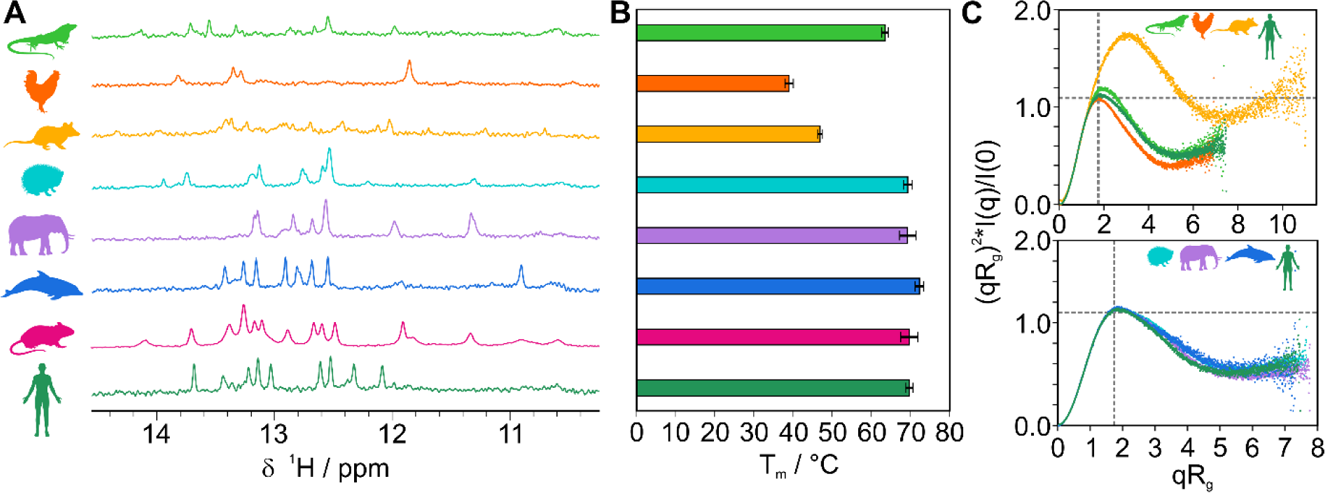
*Ox40* ADEs from multiple species form stable structures. **A)** Imino ^1^H spectra of *Ox40* species ADEs (from top to bottom: lizard, chicken, opossum, hedgehog, elephant, dolphin, mouse, and human). **B)** CD-derived melting points of ADEs aligned and color coded as in panel A). Melting points are given as average from triplicates with standard deviations (error bars). **C)** Dimensionless Kratky plots derived from SAXS measurements of ADEs. A curve maximum at the intersection of the two grey dotted lines corresponds to a globular (compact) fold. Curves with a shift to the upper right indicate partially unfolded structures (e.g. as observed for opossum).

We next used small-angle X-ray scattering (SAXS) to probe the three-dimensional shape of the ADEs (**Figure 2C** and **Supplementary Table 2**)^48^. Dimensionless Kratky plots allow to assess the fold of an RNA independent of its molecular weight^49, 50^. A bell-shaped curve maximum at the so-called Guinier-Kratky-point (i.e. the intersection of the two dashed lines in **Figure 2C**) indicates a globular fold, while shifts in the maximum to the upper right point at elongated or partially unstructured molecules. For hedgehog, dolphin, elephant and human we observe the curve maximum close to the intersection, pointing at a globular and similar fold between these four ADEs (**Figure 2C bottom panel**). This agrees well with SAXS-based tertiary structure models (**Supplementary Figure 5C**) that show a high degree of similarity. In contrast, the putative chicken and lizard ADE curves show a visible deviation from the mammalian ones (**Figure 2C top panel**). This suggests a more opened conformation for the lizard, in line with weak and broad imino proton signals in our NMR spectra. The chicken ADE shows a more bell-shaped curve, which can be explained by the formation of a partial duplex stem and the large loop. Our NMR spectra indeed confirm the existence of only four stable base pairs. This renders the overall shape more globular than the slightly extended form of the mammalian ADEs (**Supplementary Figure 1D** and **Supplementary Figure 5C**). Our results show that the *Ox40* ADE forms a stable moiety in mammals, and that this fold is robust across a broad range of temperature, which allows formation of hairpin structures at physiological conditions. Analyses of reptile and bird RNAs suggested that ADEs could have evolved from less stable and only transiently structured hairpins.

### Roquin developed a high affinity for mammalian ADEs

We wondered whether Roquin binds all species ADEs and how subtle differences in RNA sequence and shape would affect binding. Therefore, we performed gel shift assays (EMSA) of ADEs with murine/human coreROQ and extROQ (**Figure 3** and **Supplementary Figure 6**). For mammalian ADEs both protein constructs yielded discrete complex bands, confirming RNP formation (**Figure 3A**). For lizard and chicken ADEs, micromolar protein concentrations caused a smear in the gel, which suggests formation of a less stable complex that partially dissociates within the gel matrix. However, for extROQ binding occurred at lower protein concentrations (lizard) or led to formation of complex bands (chicken), albeit less sharp than for mammalian ADEs. Quantification of EMSAs (**Figure 3B and C** and **Table 2**) confirmed that Roquin binds mammalian ADEs with nanomolar affinity, while putative avian and reptilian ADEs are bound with ∼10x lower affinity (micromolar). Interestingly, elephant and dolphin ADEs showed an intermediate high-nanomolar affinity. These findings suggest that ADE binding at relevant protein concentrations only occurred in mammals, but binding was further enhanced throughout development of individual mammalian species. A comparison of K_D_ values from both proteins (**Figure 3C and D**) revealed a higher affinity of coreROQ compared to extROQ for mammalian ADEs, except for the murine one, which bound both proteins equally well. The putative lizard ADE had a higher affinity for extROQ. This can point at an accessory function of the Roquin B-site in early ADE development, where the additional RNA binding interface could facilitate RNP formation through charges in lack of a stable hairpin.

**Figure 3.**
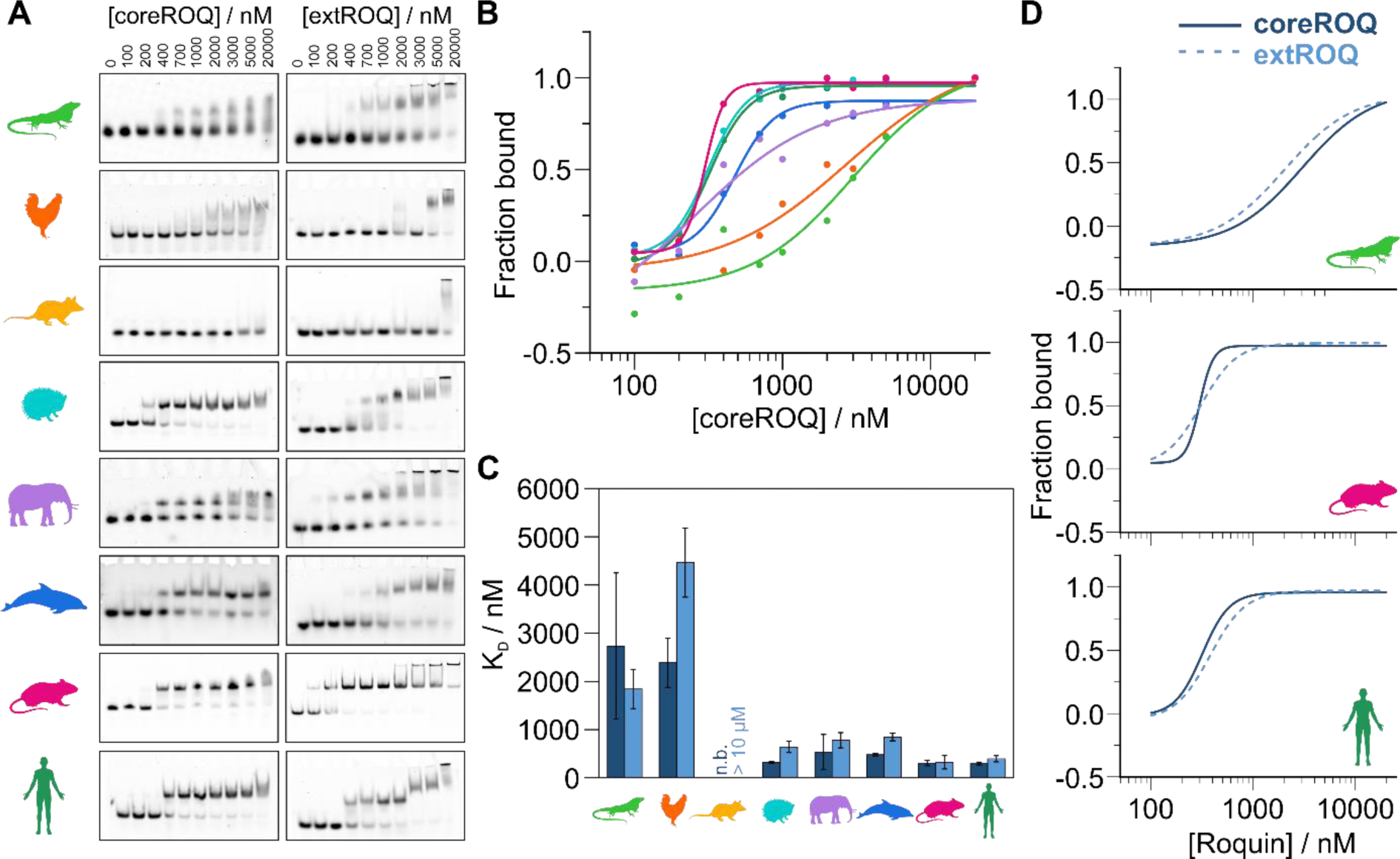
Affinity of Roquin for ADEs increased during evolution. **A)** Representative EMSAs of coreROQ (left column) and extROQ (right column) with species ADEs. Protein concentrations are given on top. Triplicates are shown in Supplementary Figure 6. **B)** CoreROQ binding curves derived from EMSAs. Plotted are averaged curves from triplicates. **C)** K_D_ values of coreROQ (dark blue) and extROQ (light blue) for species ADEs as obtained from EMSAs shown as bar plot. K_D_ values are averages from triplicates and errors are standard deviations (see Table 2). n.b. = no binding **D)** Comparison of coreROQ (dark blue, solid line) and extROQ (light blue, dashed line) binding curves with lizard, mouse and human ADEs.

**Table 2:**
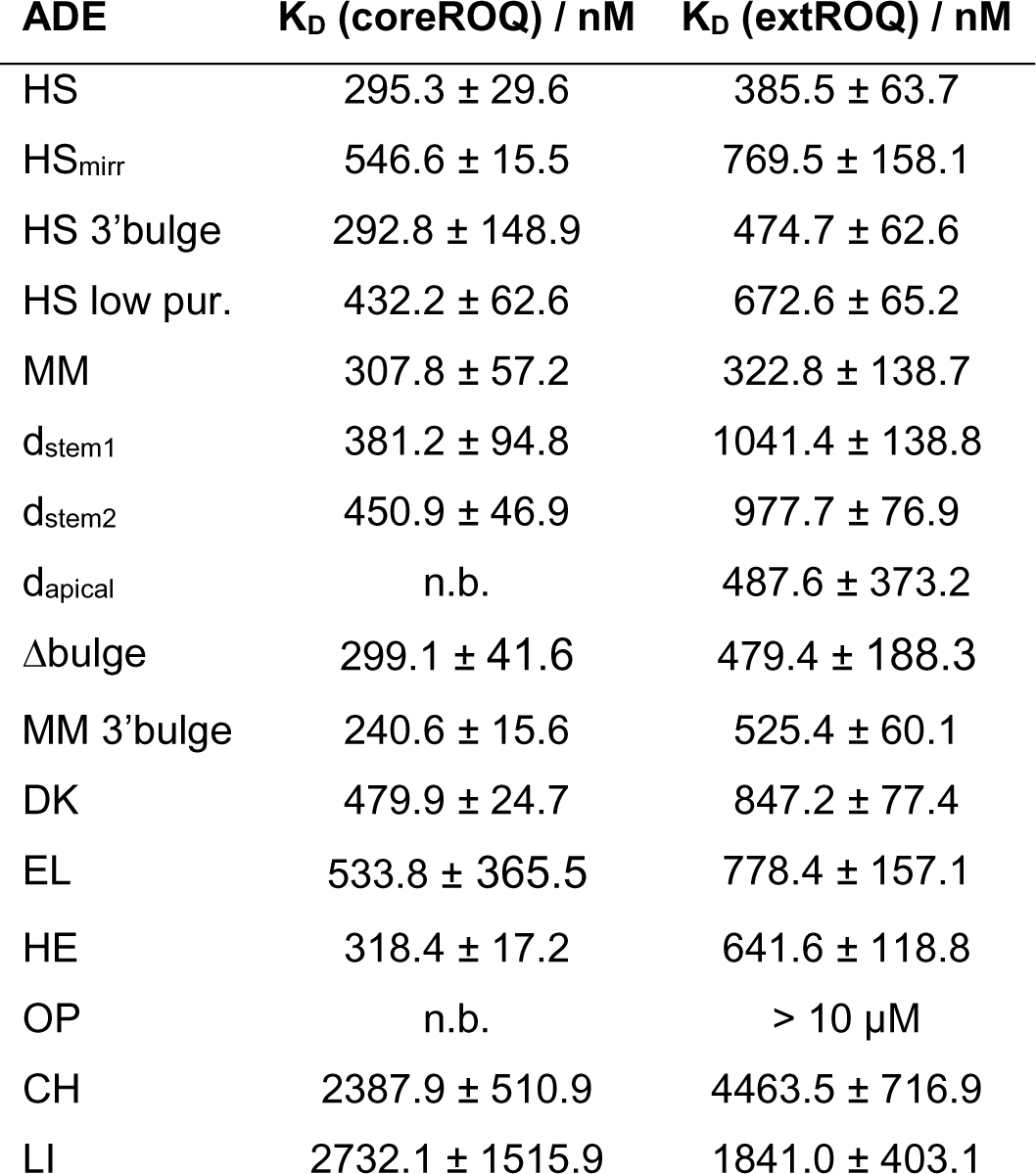
Affinities of coreROQ and extROQ for ADE RNAs. KD values are averages of n = 3; errors are standard deviations.

| ADE | $K_D$ (coreROQ) / nM | $K_D$ (extROQ) / nM |
| --- | --- | --- |
| HS | $295.3 \pm 29.6$ | $385.5 \pm 63.7$ |
| HS<sub>mirr</sub> | $546.6 \pm 15.5$ | $769.5 \pm 158.1$ |
| HS 3'bulge | $292.8 \pm 148.9$ | $474.7 \pm 62.6$ |
| HS low pur. | $432.2 \pm 62.6$ | $672.6 \pm 65.2$ |
| MM | $307.8 \pm 57.2$ | $322.8 \pm 138.7$ |
| d<sub>stem1</sub> | $381.2 \pm 94.8$ | $1041.4 \pm 138.8$ |
| d<sub>stem2</sub> | $450.9 \pm 46.9$ | $977.7 \pm 76.9$ |
| d<sub>apical</sub> | n.b. | $487.6 \pm 373.2$ |
| Δbulge | $299.1 \pm 41.6$ | $479.4 \pm 188.3$ |
| MM 3'bulge | $240.6 \pm 15.6$ | $525.4 \pm 60.1$ |
| DK | $479.9 \pm 24.7$ | $847.2 \pm 77.4$ |
| EL | $533.8 \pm 365.5$ | $778.4 \pm 157.1$ |
| HE | $318.4 \pm 17.2$ | $641.6 \pm 118.8$ |
| OP | n.b. | > 10 μM |
| CH | $2387.9 \pm 510.9$ | $4463.5 \pm 716.9$ |
| LI | $2732.1 \pm 1515.9$ | $1841.0 \pm 403.1$ |

### The ADE binding mode changed during evolution

We next set out to determine when in evolution ADEs started to be bound in the known ‘ADE-like’ fashion^24^. To monitor changes in binding on the atomic level we recorded ^1^H,^15^N-HSQC fingerprint spectra of the murine/human coreROQ protein alone (apo) and in complex with species ADEs (**Figure 4** and **Supplementary Figure 7**). Including earlier work^24, 26^, the human and murine ADEs were found to show the same pattern of chemical shift perturbations (CSPs) (**Figure 4A**). Spectra of the mammalian ROQ-ADE complexes (**Supplementary Figure 7**) show near-complete overlap with each other with only minor deviations in CSPs, suggesting a conserved binding mode. However, avian and reptilian ADEs show large differences in CSPs between each other and compared to mammals (**Figure 4B** and **Supplementary Figure 7**). This correlates well with the observed differences between these two clades and mammals with respect to predicted structure, stability and affinity.

**Figure 4.**
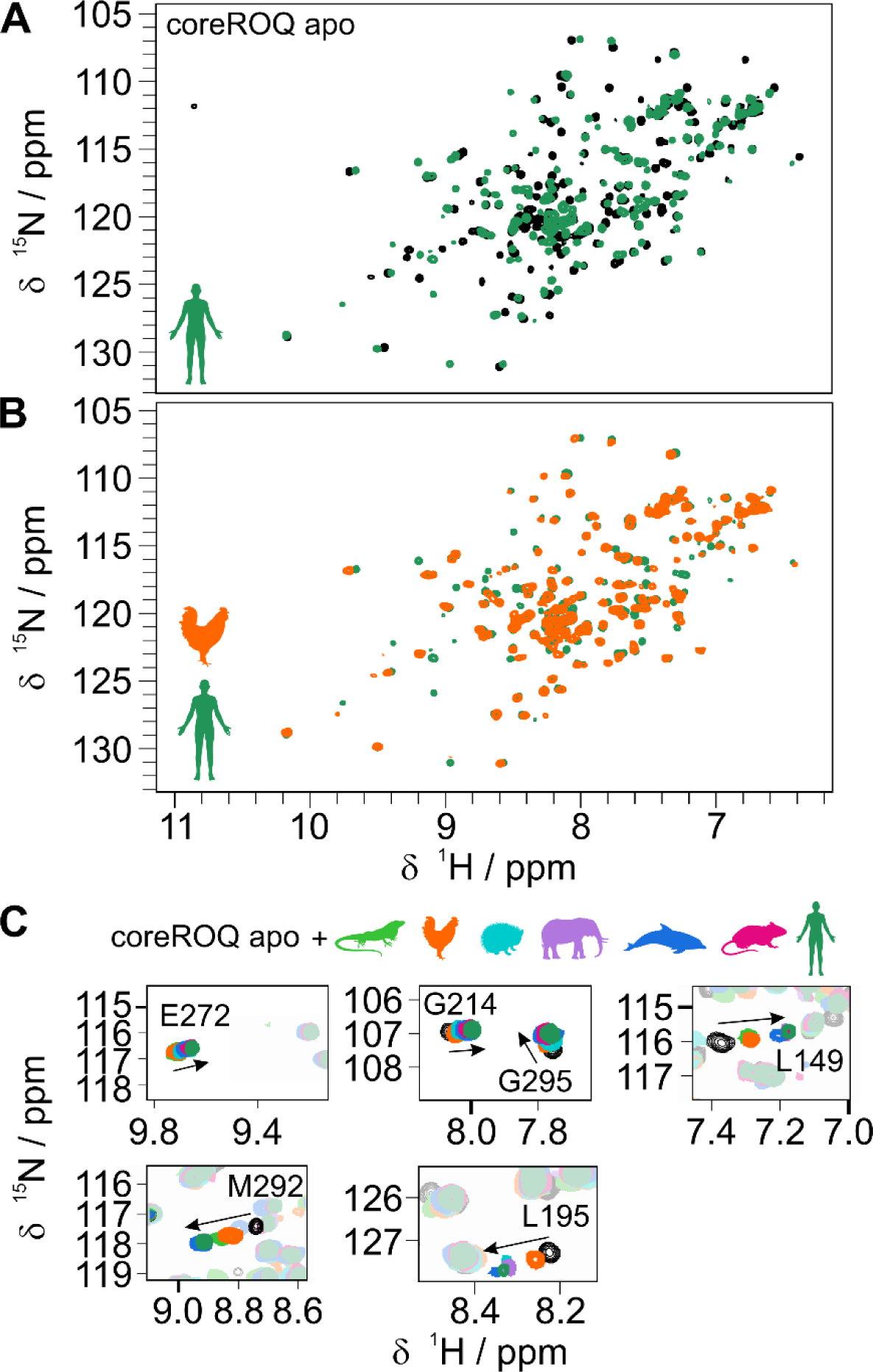
Evolution of the conserved Roquin ADE binding mode. **A)** ^1^H,^15^N-HSQC spectra of apo coreROQ (black) and in complex with the human *Ox40* ADE (green). **B)** ^1^H,^15^N-HSQC spectra of coreROQ in complex with the putative chicken (orange) or human (green) ADE. For all species comparisons refer to Supplementary Figure 7. **C)** Zoom-ins of ^1^H,^15^N-HSQC overlays of apo coreROQ (black) and in presence of species ADEs (color).

Tracking of individual amino acids in overlays of all species complexes allowed to monitor changes in CSPs along evolution: Several residues showed increased CSPs for mammalian ADEs compared to chicken and lizard ADEs (e.g. L149, M292; **Figure 4C**). Others, e.g. L19, exhibited stronger line broadening for chicken and lizard, which is often indicative of changes in affinity. Together, our NMR data capture and support the gradual changes in affinity throughout ADE evolution and recapitulate the formation of a conserved ADE binding mode.

### Stem stability affects Roquin binding

From our CD and EMSA experiments we conclude that formation of a stable stem-loop structure was a key step in evolution to generate high-affinity interactions between RNA and Roquin. We hence tested murine ADE variants with destabilizing stem mutants for protein binding (**Figure 5**). Mutations of GC base pairs to AU base pairs in different stem regions (**Figure 5A**) showed a shift in CD melting points to lower temperatures (**Figure 5B**), confirming the destabilizing effect of the mutations. Additionally, we deleted the central bulge (**Figure 5A**), which stabilized the ADE as observed in an increased melting point (**Figure 5B**). Gel-shift assays of the murine ADE variants with core and extROQ (**Figure 5C** and **Supplementary Figure 8**) showed formation of complex bands. However, no band shift was observed for coreROQ incubated with the ADE comprising destabilizing mutations in the apical stem (d_apical_). This confirms that the apical stem-loop is the only binding site for coreROQ and that a strong reduction in stem stability severely perturbs RNP formation through the Roquin A-site. In extROQ the B-site partially compensates this effect as assessed by complex band formation. Quantification of EMSAs (**Figure 5D**) revealed that destabilization of the central stem regions (d_stem1_ and d_stem2_) had only a minor effect on coreROQ binding. On the contrary, extROQ engagement with these ADE variants was significantly perturbed and led to a decrease in affinity of ∼3x (**Table 2**). Interestingly, K_D_ values of both coreROQ and extROQ for the Δbulge ADE variant were almost not altered from the wildtype (**Figure 5D**). A slight difference for extROQ, which is within the error, could point at a structure-mediated effect recognized through the B-site. Despite this small effect, further stabilization of the wildtype ADE stability does neither enhance nor reduce the affinity of Roquin.

**Figure 5.**
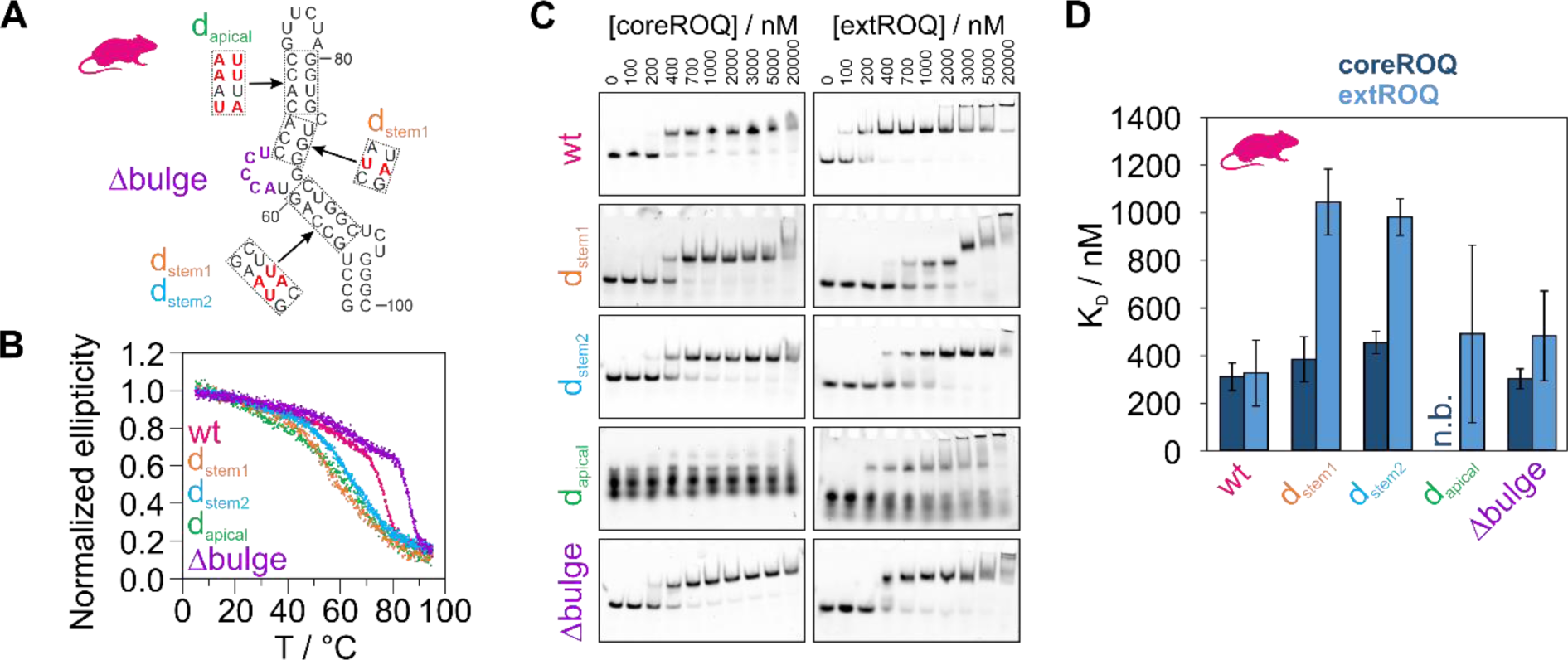
ADE stability mutants modulate Roquin binding affinity. **A)** Secondary structure of murine *Ox40* ADE wt. Red nucleotides in dashed boxes indicate sites of mutations. The corresponding constructs are denoted besides the boxes. Nucleotides deleted in a Δbulge version are colored in purple. **B)** Normalized CD melting curves of ADE variants shown in A). **C)** Representative EMSAs of coreROQ (left column) and extROQ (right column) with stability variants of the murine *Ox40* ADE from A). Protein concentrations are given on top. Triplicates are shown in Supplementary Figure 8. **D)** K_D_ values of coreROQ (dark blue) and extROQ (light blue) for ADE stability variants as obtained from EMSAs shown as bar plot. K_D_ values are averages from triplicates and errors are standard deviations (see Table 2). n.b. = no binding

### RNA determinants required for high-affinity Roquin binding

Changes in stem stability do not explain the increase in affinity throughout mammalian development (**Figure 3C**), where ADEs are similar in both shape and stability (**Figure 2B** and **C**). We hence examined features of sequence and structure that differ among mammals. Interestingly, an internal bulge with an asymmetric number of unpaired nucleotides evolved directly below the upper four-to-five base pair hairpin stem (**Supplementary Figure 1D**), which harbors the known coreROQ binding site. While elephant and dolphin possess a 3’ bulge (**Supplementary Figure 1D** and **Supplementary Table 3**), hedgehog, human and mouse ADEs form a 5’ bulge. We observed a positive correlation of affinity with the presence of a 5’ bulge (**Figure 6A**) for extROQ. Binding affinity also correlated well with the size of the bulge. Interestingly, binding affinity of coreROQ to ADEs did not correlate with a 5’ or 3’ bulge, confirming no interaction of coreROQ with this region of the ADE hairpin (**Figure 6A**). We thus hypothesized that bulge inclusion in either stem half and the bulge size make up a structural feature that is read out by Roquin, in particular by the B-site. We tested our hypothesis with two technical ADE mutants: For the human and murine ADE we swapped the bulge to the 3’ stem (**Supplementary Figure 9A**) and analyzed Roquin binding by EMSAs (**Figure 6B** and **Supplementary Figure 9B**). For the human ADE, we observed no change in affinity for coreROQ (**Figure 6C**), while the murine ADE 3’ bulge variant even showed a small increase in affinity. Affinity of extROQ showed a small decrease for the human 3’ bulge ADE compared to the wildtype ADE, while the reduction in affinity by a factor of ∼1.6x was larger for the murine version (**Figure 6C**). We suggest the different responsiveness between the two species is caused by the increased size of the bulge for the mouse (**Figure 6A**).

**Figure 6.**
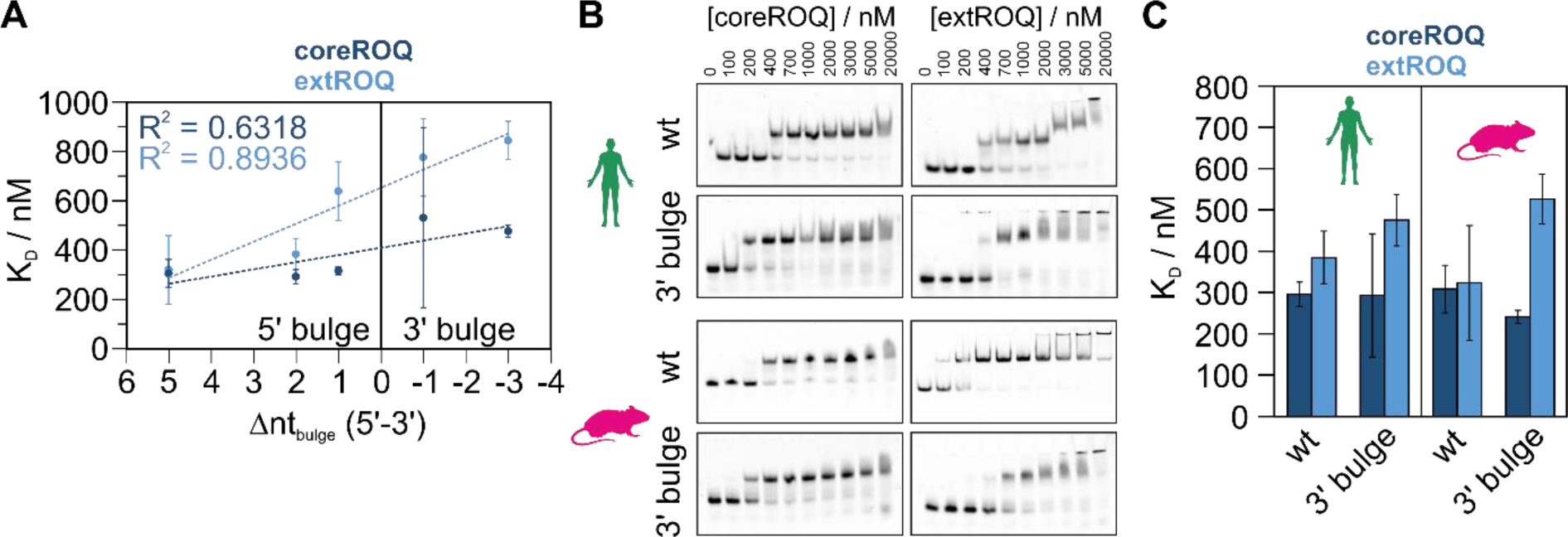
The central ADE bulge affects Roquin binding affinity. **A)** Correlation plot of coreROQ (dark blue) and extROQ (light blue) affinities for mammalian ADEs (taken from Figure 3C) with the number of unpaired nucleotides in the central bulge of the respective ADE (see Supplementary Table 3). Note that the difference between 5’ and 3’ unpaired nucleotides is plotted. Positive values thus indicate a bulge within the 5’ stem and negative values within the 3’ stem. **B)** Representative EMSAs of coreROQ (left column) and extROQ (right column) with human and murine wt ADEs and a version with a 3’ bulge (see Supplementary Figure 9). Protein concentrations are given on top. Triplicates are shown in Supplementary Figure 9. Wt EMSAs are the same as in Figure 3A. **C)** K_D_ values of coreROQ (dark blue) and extROQ (light blue) for wt and 3’ bulge ADEs as obtained from EMSAs shown as bar plot. K_D_ values are averages from triplicates and errors are standard deviations (see Table 2).

### Roquin reads out stem geometry

A close inspection of the species ADE sequences revealed a purine content in the 3’ stem of 46.2 % to 57.1 % (**Supplementary Table 4**). Binding affinity for both coreROQ and extROQ showed a good correlation with the 3’ purine content, albeit a stronger dependency was observed for extROQ binding (**Figure 7A**). To probe for purine sensitivity, we lowered the 3’ purine content of the human ADE by swapping base pairs (**Figure 7B**) from 53.8 % to 23.1 % (**Supplementary Table 4**). In EMSAs (**Figure 7C**) a low 3’ purine content led to a reduction in affinity by 1.5-fold for coreROQ and 1.7-fold for extROQ (**Figure 7D** and **Table 2**). Our data thus highlight a potential role of the central stem bulge and the purine content of the 3’ stem for ADE complex formation with Roquin.

**Figure 7.**
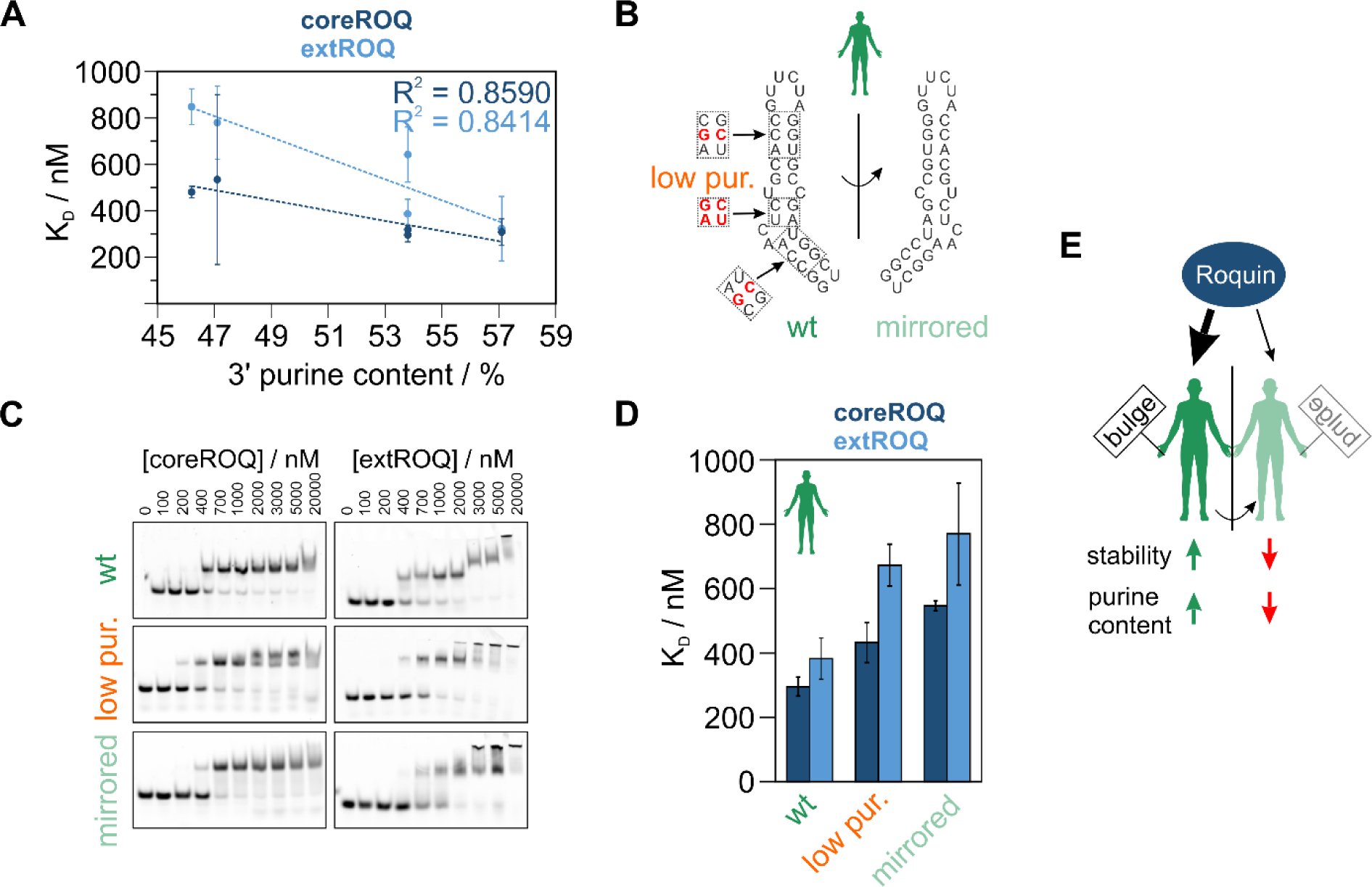
The impact of ADE geometry and purine content on Roquin binding affinity. **A)** Secondary structure of human *Ox40* ADE wt and a mirrored version. Note that the central stem is mirrored, while the basal stem and the loop orientation are maintained with respect to the wildtype. Red nucleotides in dashed boxes indicate sites of mutations within an ADE mutant with low purine content in the 3’ stem. **B)** Representative EMSAs of coreROQ (left column) and extROQ (right column) with human ADE variants from A). Protein concentrations are given on top. Triplicates are shown in Supplementary Figure 10. Wt EMSAs are the same as in Figure 3A. **C)** K_D_ values of coreROQ (dark blue) and extROQ (light blue) for human ADE variants as obtained from EMSAs shown as bar plot. K_D_ values are averages from triplicates and errors are standard deviations (see Table 2). **D)** Roquin shows a binding preference for ADEs with a bulge in the 5’ stem. Increased stability and a high purine content in the 3’ stem increase affinity. Overall, the interplay of geometric factors, stability and sequence fine tunes Roquin-ADE complex formation.

At this stage, our experiments with swapped bulges, lower purine content and changes in stem stability showed detectable yet moderate effects on binding affinity. We next speculated that the precise interplay of loop sequence, geometry and stability renders a high-affinity Roquin target. We thus created a mirrored version of the human ADE (**Figure 7B**) where only the loop and basal stem orientation was preserved and confirmed its structural integrity and the mirrored shape by NMR spectroscopy (compare **Supplementary Figure 10A** and **Supplementary Figure 3B**). EMSAs confirmed formation of stable complexes (**Figure 7C** and **Supplementary Figure 10B**), but with K_D_ values of 546 and 769 nM for core and extROQ, respectively (compared to 295 and 385 nM for the human wildtype ADE; **Table 2**). The affinity reduction for the mirrored ADE strongly indicates that Roquin recognizes a defined geometry and requires a precise interplay of loop sequence orientation and stem geometry (**Figure 7D**). However, NMR spectra confirmed a similar binding mode for both wildtype and mirrored human ADEs (**Supplementary Figure 10C**). This suggests that Roquin forces the suboptimal geometry of the mirrored ADE into a preferred fold, resulting in the lowered affinity.

## DISCUSSION

CDEs are abundant RNA *cis* elements specifically recognized by Roquin to control transcript levels of target genes. Shape-recognition assures highly specific complex formation and enables effective posttranscriptional control. ADEs are much less abundant in 3’UTRs than CDEs and structural information about ADEs and their engagement with Roquin is limited. High-resolution structures of a SELEX-derived hexaloop-stem and the *Ox40* ADE^24, 26^ confirmed a preference for a GUUUUA/GUUCUA loop sequence by Roquin, leading to comparable affinities as for CDEs (171 nM for the *Ox40* ADE, 149 nM for the *Ox40* CDE)^26^. For *NFKBID* a functionally relevant ADE has been confirmed *in vivo*^16^. Braun et al. predicted further 19 ADEs in human genes, of which three were found in proximity to a CDE^28^. Folding probabilities of these ADEs varied significantly questioning an ADE-like fold. However, these predicted ADEs^28^ as well as available experimental data on the *Ox40* ADE^26^ suggested a large structural variety within this class of RNA *cis* elements.

Our comparative analysis of *Ox40* ADEs from different species contributes to empirically filling the gap in sparse ADE data. We showed that mammalian ADEs are similar in size, shape, and stability and are bound by Roquin with nanomolar affinity, suggesting positive selection of RNA parameters during evolution by the protein. A comparison to bird and reptile analoga suggested that an increase in thermal stability and formation of a stable hairpin structure led to increased binding affinity and was a critical event in the evolution of ADEs. Despite the missing annotation of the herein presented chicken and lizard ADEs, our findings illustrate how ADEs (and other structured *cis* elements) could have evolved from AU-rich, i.e. rather unstructured RNAs. Roquin binding to these putative ADEs in the low micromolar range is comparable to Roquin ZnF domain binding to AREs^13^. As Roquin is highly conserved across vertebrates, this might point at an increasing relevance of the ROQ domain in more highly developed immune systems, while at early time points in evolution gene regulation could have relied on (low-affinity) AU-rich elements. This agrees well with the observed induction of RNA structure by Roquin in AU-rich, unstructured RNAs in *UCP* that can fold into CDEs^47^. Interestingly, in line with this, we observed that the Roquin B-site had a greater contribution to binding of these progenitor ADEs. We conclude that the B-site, which was shown to interact with dsRNA^25^, i.e. rather charge driven and sequence unspecific, could have selected for RNAs that form (transient) hairpin structures until stable RNA folds evolved for specific engagement with the coreROQ domain.

Our data expands the target spectrum of Roquin. Different from the known consensus loop motif, we show that substitutions at loop positions 1 and 4 are tolerated, and Roquin can even cope with increased loop sizes: The octaloop from chicken could be recognized in a hexaloop like fashion with a rather weak A-A closing base pair. This agrees with Roquin binding to an octaloop in the *ICOS* 3’UTR with low micromolar affinity^27^. However, a poly(U) loop background needs to be embedded in a hairpin context, as we observed no binding to the large and flexible opossum ADE loop. ADEs could thus have evolved from AU-rich, unstable elements and initially might have included larger loops.

While RNA elements are often compared across different genes, our comparison of ADEs from the same gene but of different species allows to evaluate the contribution of subtle changes in sequence and structure to complex formation. In line with previous observations^51^, we showed that high-affinity Roquin binding requires formation of a stable stem. We gave evidence that a central bulge 4 to 5 bp basal to the loop affected Roquin binding, especially via the B-site. Bulge integration into the 5’ ADE stem combined with an increase purine content in the 3’ stem enhanced interaction of the B-site with the central stem. While an increased 3’ purine content had been suggested before, not all predicted ADEs possess such a purine stack^28^. 3’ purine stacks were shown for CDEs to widen the RNA major groove and thereby facilitate Roquin engagement with the 5’ stem side^51^. This causes optimal prearrangement of the RNA structure and facilitates complex formation compared to hairpins without purine stacks. Hence, purine rich 3’ stems provide yet another layer of fine-tuning of Roquin RNPs required for a subset of targets only and are especially recognized by the Roquin B-site. In line with this, our data on *Ox40* species ADEs and predicted ADEs showed that a central bulge is no absolute requirement for Roquin binding, despite its positive effect on affinity. Together, we confirmed that the Roquin B-site prefers duplex RNA regions^25^, but it is sensitive to geometric deviations from a perfect duplex and can readout geometric parameters, e.g. mirrored shapes. While the coreROQ domain has a well-defined target consensus for a stable apical stem-loop, which our data corroborate, the B-site is more promiscuous with respect to sequence and structure. We conclude that the precise interplay of Roquin A and B-site integrates high specificity for stem-loops with a platform to readout RNA stem stability and geometry. For a high-affinity interaction, the concerted recognition of these defined apical and central stem elements is crucial.

Altogether, our study provides insight into the evolution of ADE stem-loop features from potential progenitor ADEs, which differ significantly from the known ADE consensus forming stable hairpin structures. This development of RNA *cis* element structure is accompanied by an increase in Roquin binding affinity. The interaction of Roquin with ADEs might not have existed early in evolution but could have rather evolved as a second RNP besides the Roquin-CDE complex. The evolutionary snapshots in our study provide us detailed information on key steps of the evolution of RNA determinants for an optimized, highly adapted RNP in coincidence with the development of the adaptive immune system.

## SUPPLEMENTARY DATA

Supplementary Data are available at XXX online.

## ACKNOWLEDGEMENT

We acknowledge excellent technical support by Katharina Targaczewski and thank the team of SAXS at Beamline P12, DESY Hamburg for measurement support.

