## Supplementary material for "Evolution of the RNA alternative decay *cis* element into a high-affinity target for the immunomodulatory protein Roquin": Tants_etal_Supplement.pdf

### SUPPLEMENTARY TABLES

**Supplementary Table 1:** Melting points of species ADEs as determined by CD. $T_m$ values are averages from triplicates and standard deviations are given as errors.

| **ADE** | **$T_m$ / °C** |
|---|---|
| LI | 63.48 ± 0.83 |
| CH | 38.95 ± 1.08 |
| OP | 46.85 ± 0.54 |
| HE | 69.32 ± 1.11 |
| EL | 69.26 ± 2.10 |
| DK | 72.33 ± 1.08 |
| MM* | 69.78 ± 2.25 |
| HS | 69.69 ± 0.89 |

*Note that the murine ADE is taken from[26]. The difference in $T_m$ can be explained by the different buffer systems used.

**Supplementary Table 2:** $R_g$ of species ADEs as determined by SAXS. Concentrations used for measurements are given. For measurements with $MgCl_2$, 2 mg/ml of RNA were used.

| **ADE** | **Rg / Å** | | |
|---|---|---|---|
| | 2 mg/ml | 3 mg/ml | +1 mM $MgCl_2$ |
| LI | 17.06 ± 0.82 | 16.58 ± 0.10 | 16.19 ± 0.03 |
| CH | 16.50 ± 0.65 | 15.58 ± 0.04 | 15.97 ± 0.02 |
| OP | 24.17 ± 0.17 | 24.89 ± 0.23 | 23.67 ± 0.18 |
| HE | - | 17.28 ± 0.10 | 18.02 ± 0.02 |
| EL | 18.32 ± 0.03 | 17.51 ± 0.05 | 18.17 ± 0.02 |
| DK | - | 16.50 ± 0.03 | 16.46 ± 0.02 |
| MM* | 20.84 ± 0.03 | 25.36 ± 0.17 | - |
| HS | 18.85 ± 0.13 | 16.83 ± 0.03 | 19.74 ± 0.11 |

*Note that the murine ADE is taken from[26] and significantly larger, which explains the increased $R_g$.

**Supplementary Table 3:** Distribution of unpaired nucleotides within the central bulge of mammalian ADEs. nt = nucleotides

| ADE | Unpaired 5' nt | Unpaired 3' nt | Difference 5'-3' |
|---|---|---|---|
| HS* | 2 (5) | 0 (3) | 2 (2) |
| $HS_{mirr}$ | 0 (3) | 2 (5) | -2 (-2) |
| HS 3' bulge | 0 (3) | 2 (5) | -2 (-2) |
| MM | 5 | 0 | 5 |
| MM 3'bulge | 0 | 5 | -5 |
| DK | 1 | 4 | -3 |
| EL | 3 | 4 | -1 |
| HE | 4 | 3 | 1 |

*Values in parentheses assume opening of both bulges as indicated in Supplementary Figure 3B based on $^{1}H$,$^{1}H$-NOESY secondary structure assignment. Note that both secondary structures show the same distribution of unpaired nucleotides.

**Supplementary Table 4:** Purine content of ADE stem halves in percent.

| ADE | 5' purines | 3' purines |
|---|---|---|
| HS | 40.0 | 53.8 |
| $HS_{lowpur}$ | 66.7 | 23.1 |
| MM* | 31.8 | 57.1 (47.8) |
| DK | 30.0 | 46.2 |
| EL | 21.4 | 47.1 |
| HE | 26.7 | 53.8 |

*Note that for the mouse only stem residues 29-42 were considered, to compensate for the larger structure. The value in parentheses gives the full 3' stem purine content.

### SUPPLEMENTARY FIGURES

#### Supplementary Figure 1:

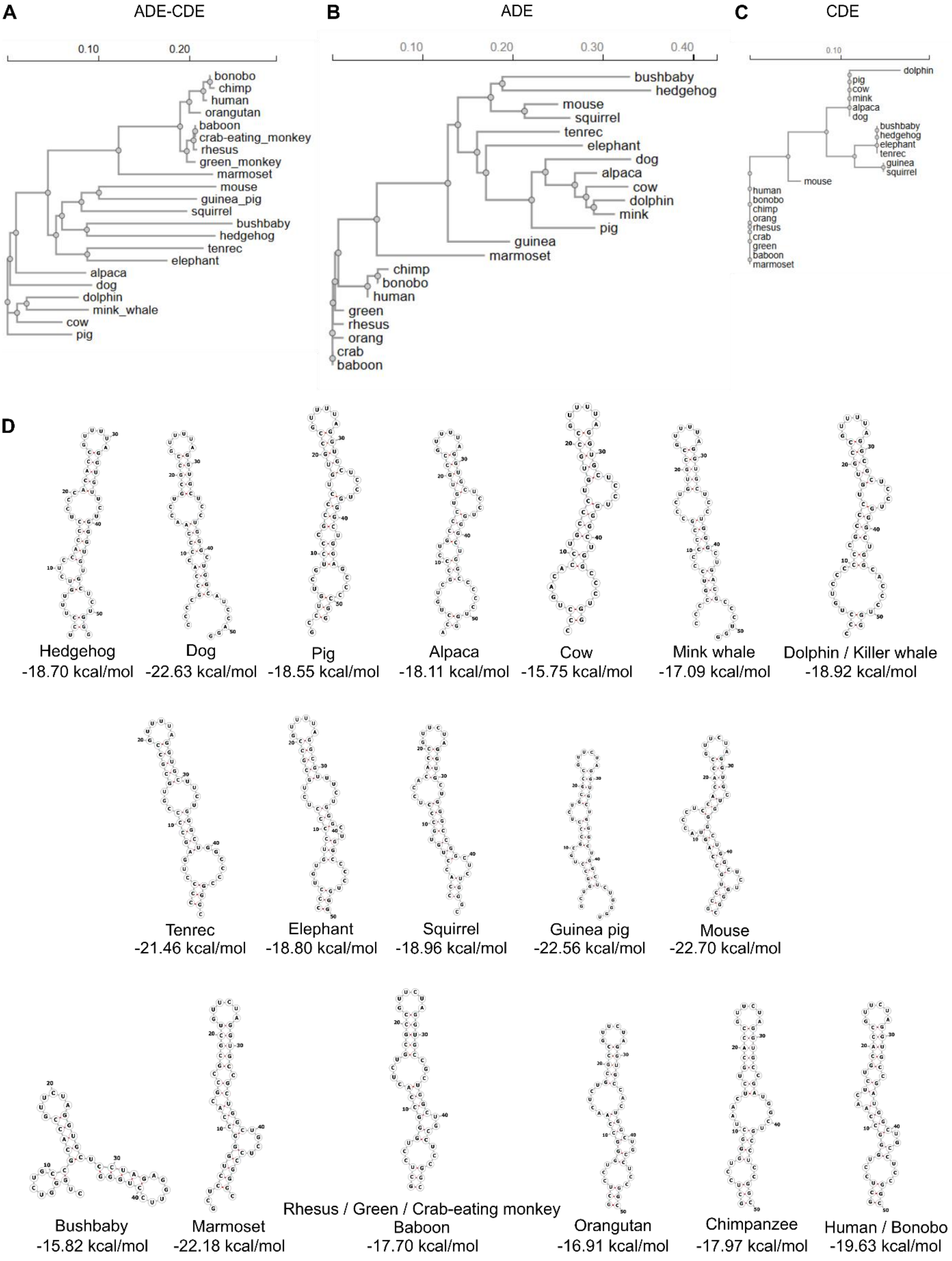


**Supplementary Figure 1.** Conservation of *Ox40* ADE-CDE cassette among vertebrates. **A)** Phylogenetic tree of tandem ADE-CDE cassette based on sequence alignment from Figure 1A. **B)** and **C)** phylogenetic trees of ADE and CDE, respectively, based on sequence alignment from Figure 1A.

**Supplementary Figure 2:**

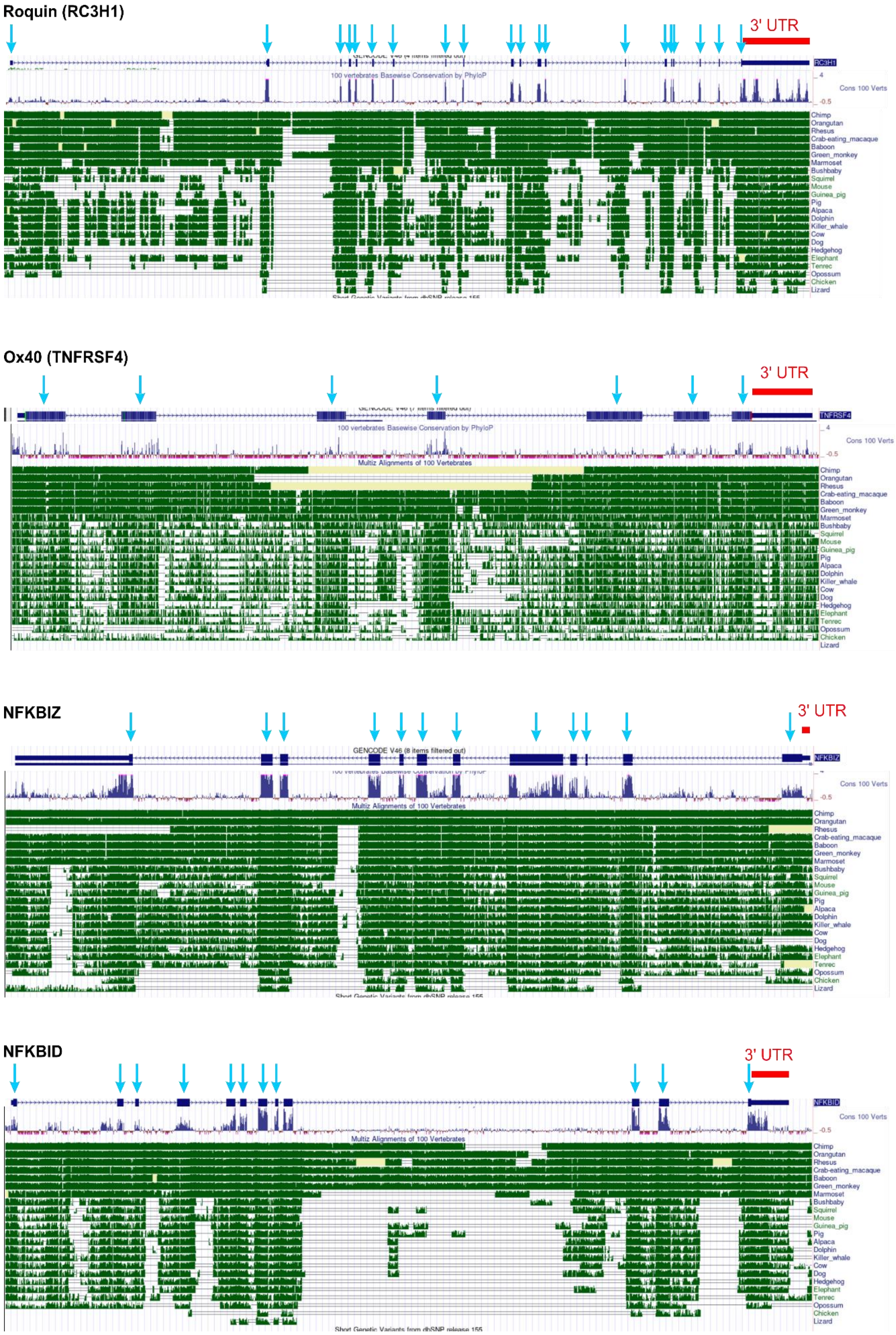


**Supplementary Figure 2.** MultiZAlignment of Roquin and targets of Roquin from 100 vertebrates. Exons within the pre-mRNA are highlighted by blue arrows, 3’UTRs by red bars. Below the mRNA scheme the conservation scores from an alignment of 100 vertebrates is shown. In green conserved regions of selected species used in Figure 1A are depicted. The panels show the conservation of **A)** Roquin (*RC3H1*), **B)** *Ox40*, **C)** *NFKBIZ* and **D)** *NFKBID*.

**Supplementary Figure 3:**

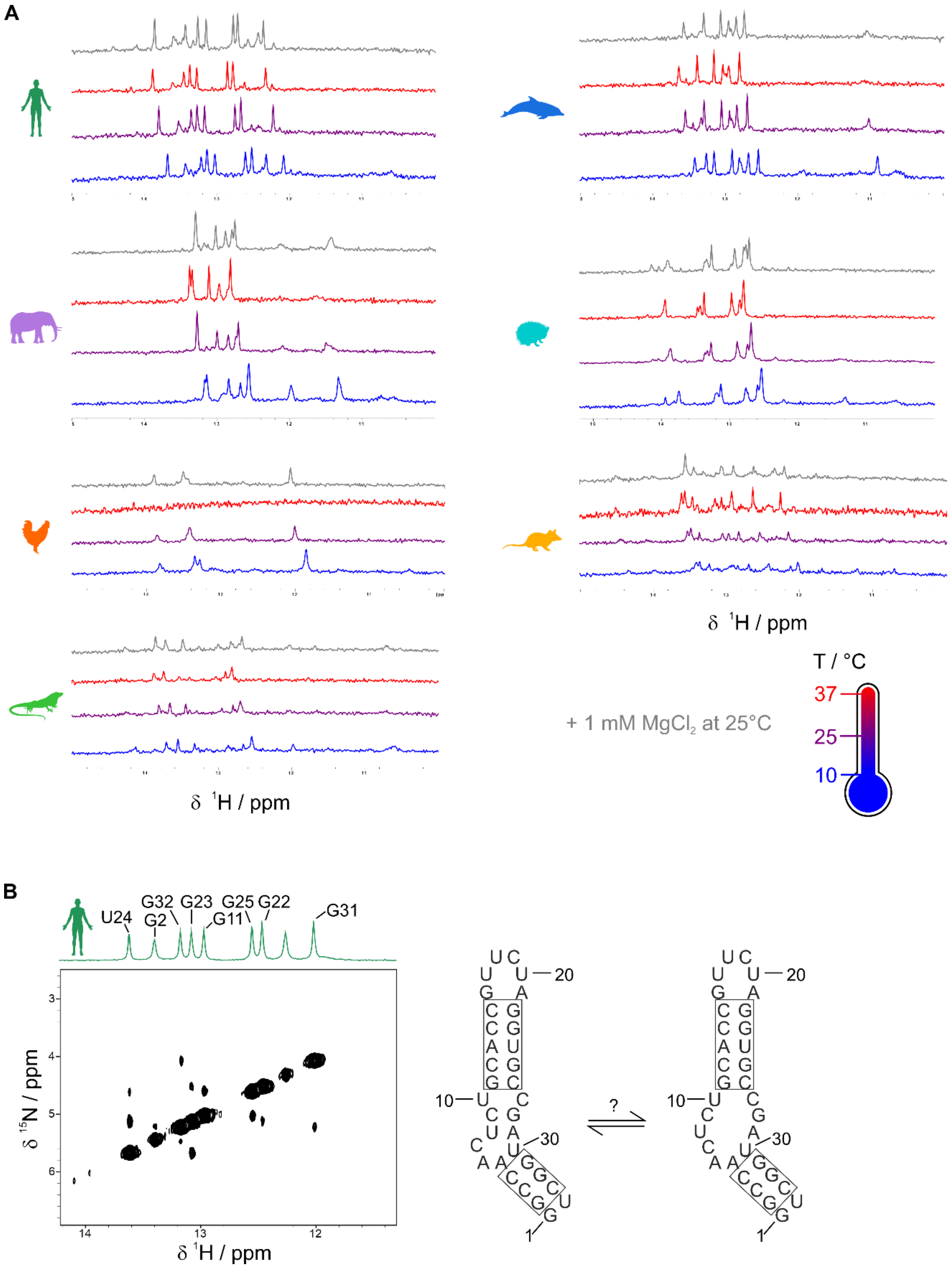


**Supplementary Figure 3.** NMR spectroscopy confirms stable fold of *Ox40* ADEs. **A)** For each species indicated by icons imino $^1$H spectra at 10, 25 and 37°C are shown (blue, purple, and red, respectively). On top in grey, the corresponding imino $^1$H spectrum of the ADE in presence of 1 mM $MgCl_2$ at 25°C is shown. **B)** $^1$H,$^1$H-NOESY spectrum of the human ADE. Imino proton resonance assignment is shown in the corresponding 1D spectrum on top. Confirmed base pairs are boxed in the secondary structure prediction on the right. Based on the assignment, a second conformation could form, with a partially opened bulge.

**Supplementary Figure 4:**

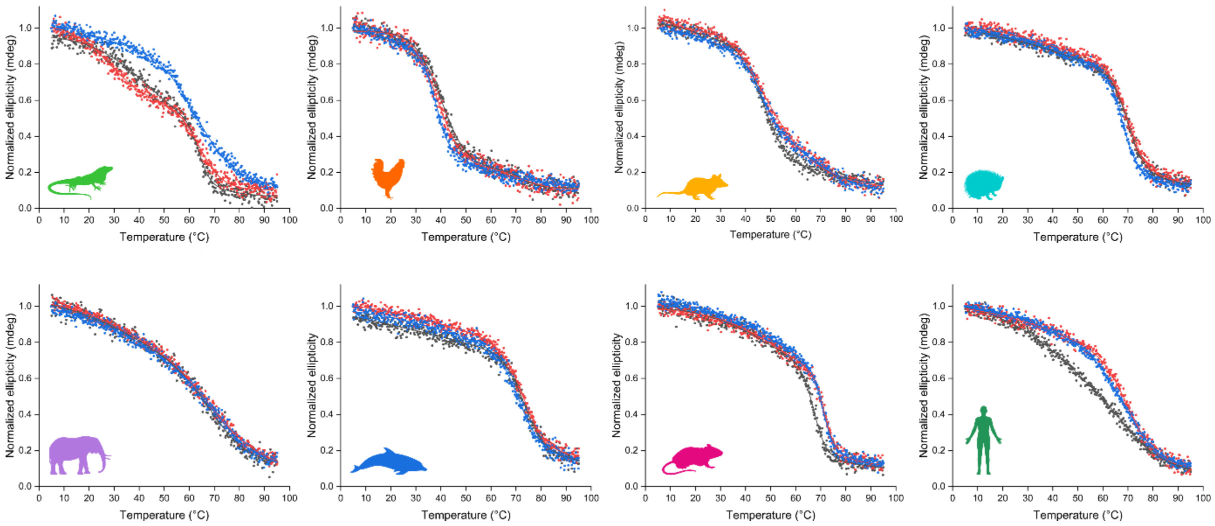


**Supplementary Figure 4.** CD melting curves of species ADEs as indicated by icons. Shown are triplicates (black, blue and red curve). Averaged melting points are plotted in Figure 2B.

**Supplementary Figure 5:**

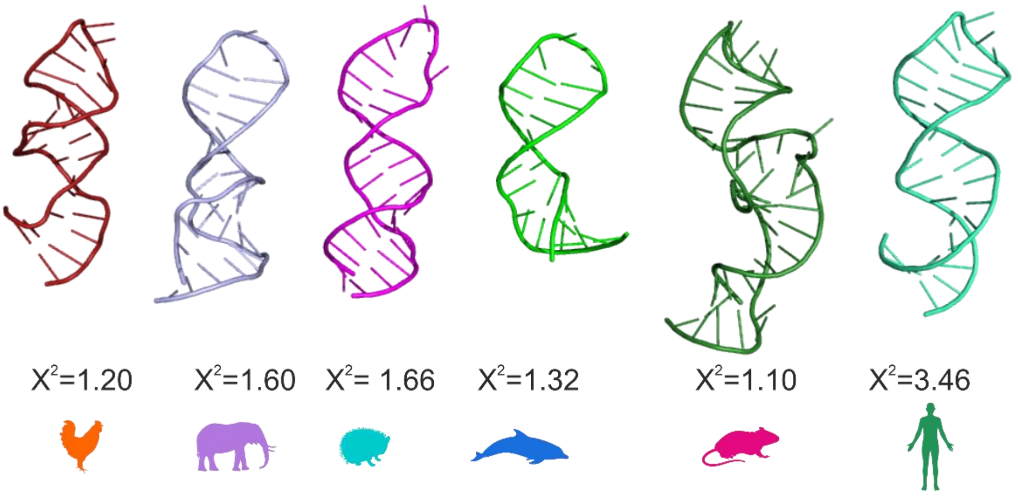


**Supplementary Figure 5.** SAXS-based RNA Masonry models of *Ox40* ADEs. Below $X^2$ values are given for the fit quality of the experimental and a theoretical SAXS curve. Note that the murine ADE model is taken from[26].

**Supplementary Figure 6:**

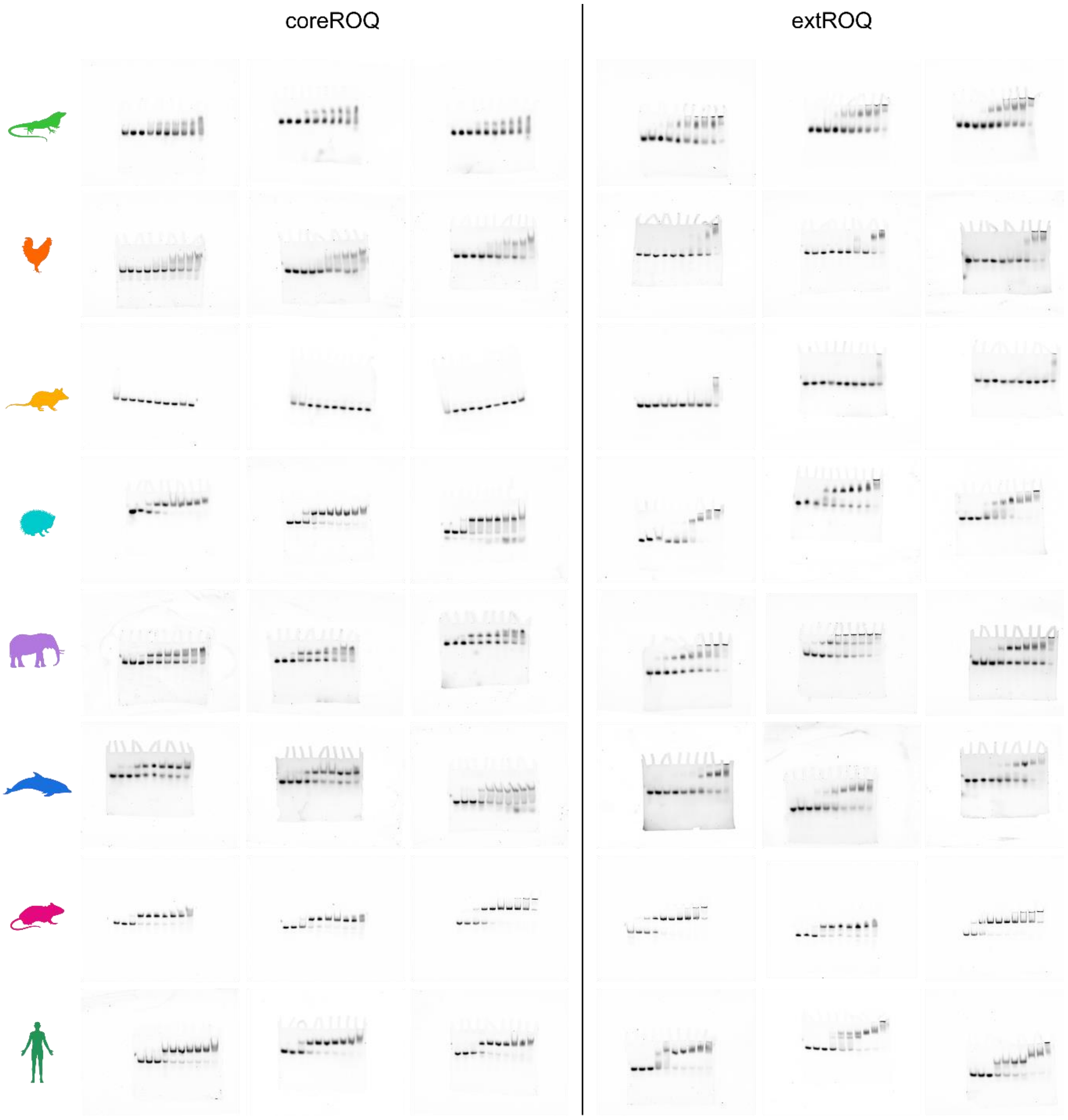


**Supplementary Figure 6.** EMSAs of coreROQ (left) and extROQ (right) with species ADEs. Shown are triplicates. Calculated $K_D$ values are averages and plotted in Figure 3C and listed in Table 2.

**Supplementary Figure 7:**

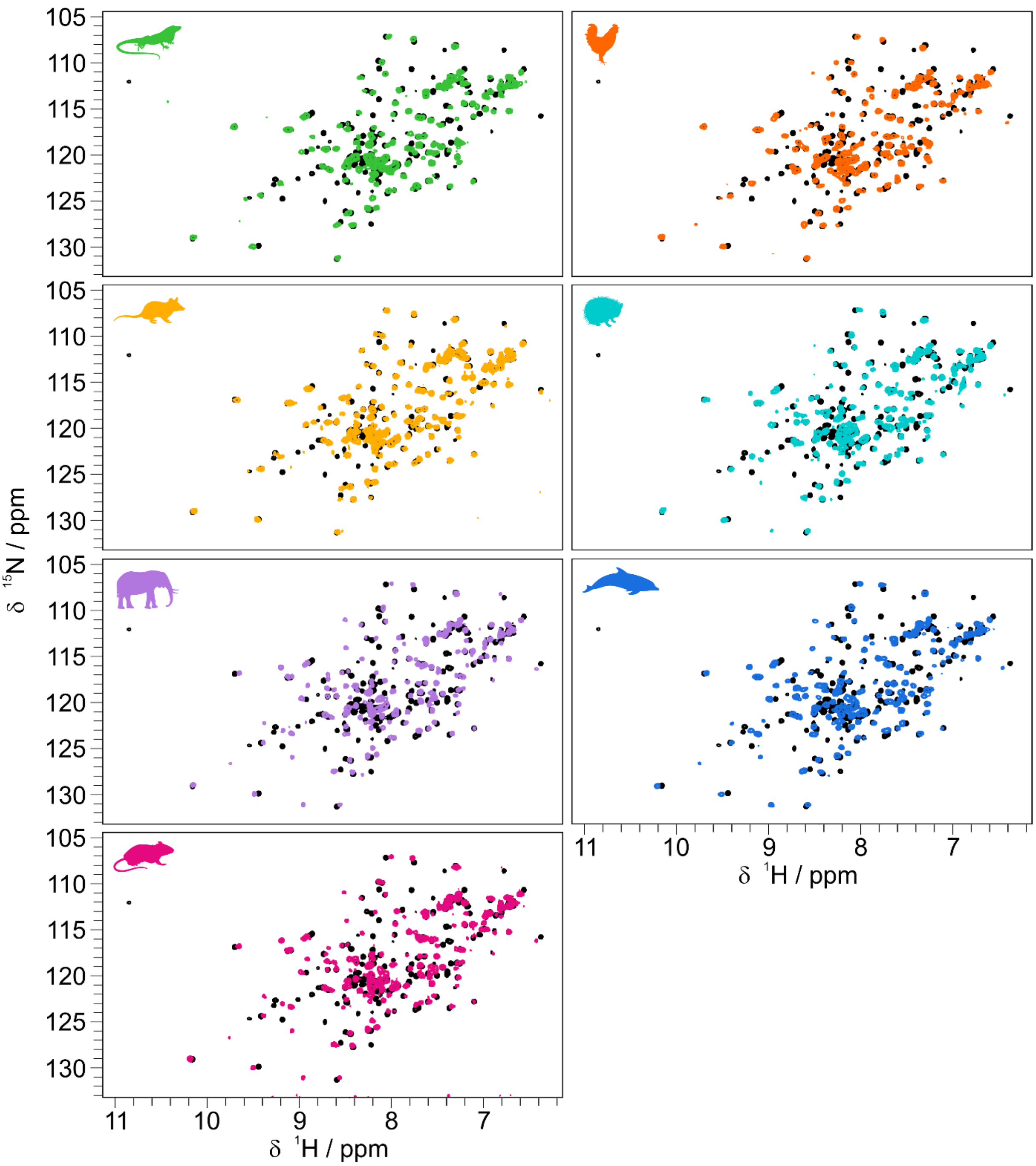


**Supplementary Figure 7.** $^{1}H,^{15}N$-HSQC overlays of apo coreROQ (black) and species ADEs (color).

**Supplementary Figure 8:**

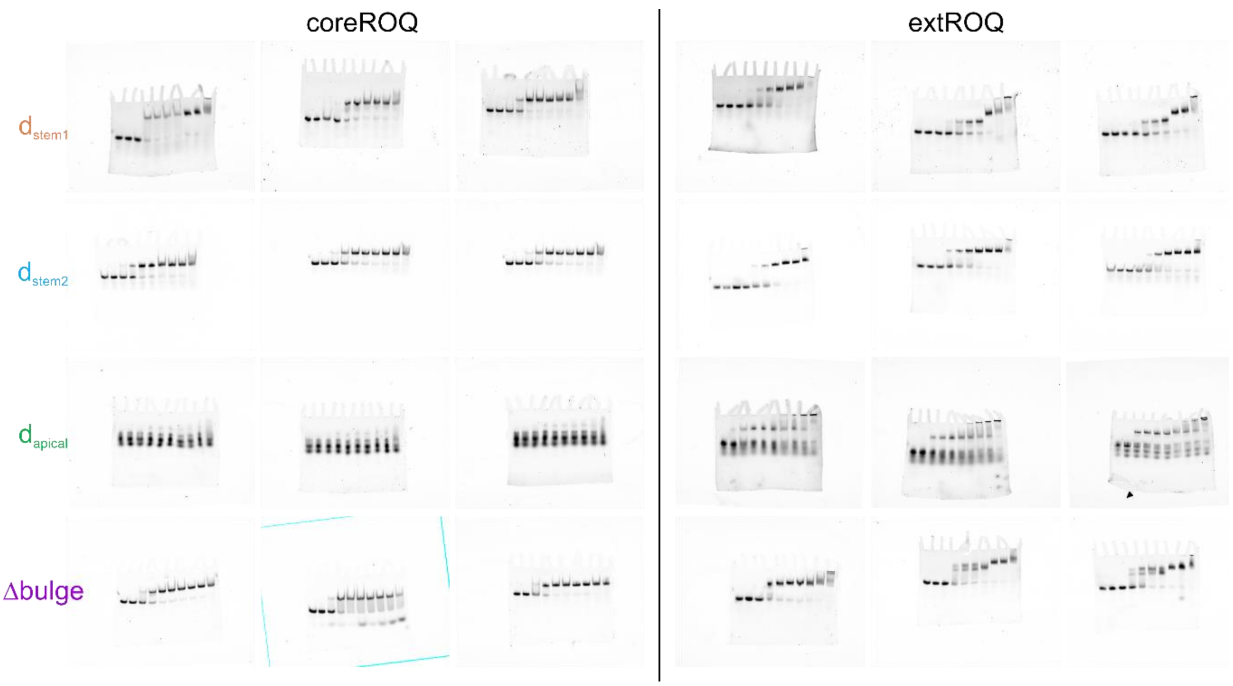


**Supplementary Figure 8.** EMSAs of coreROQ (left) and extROQ (right) with destabilized murine ADEs and a Δbulge variant. Shown are triplicates. Calculated $K_D$ values are averages and plotted in Figure 5D and listed in Table 2.

**Supplementary Figure 9:**

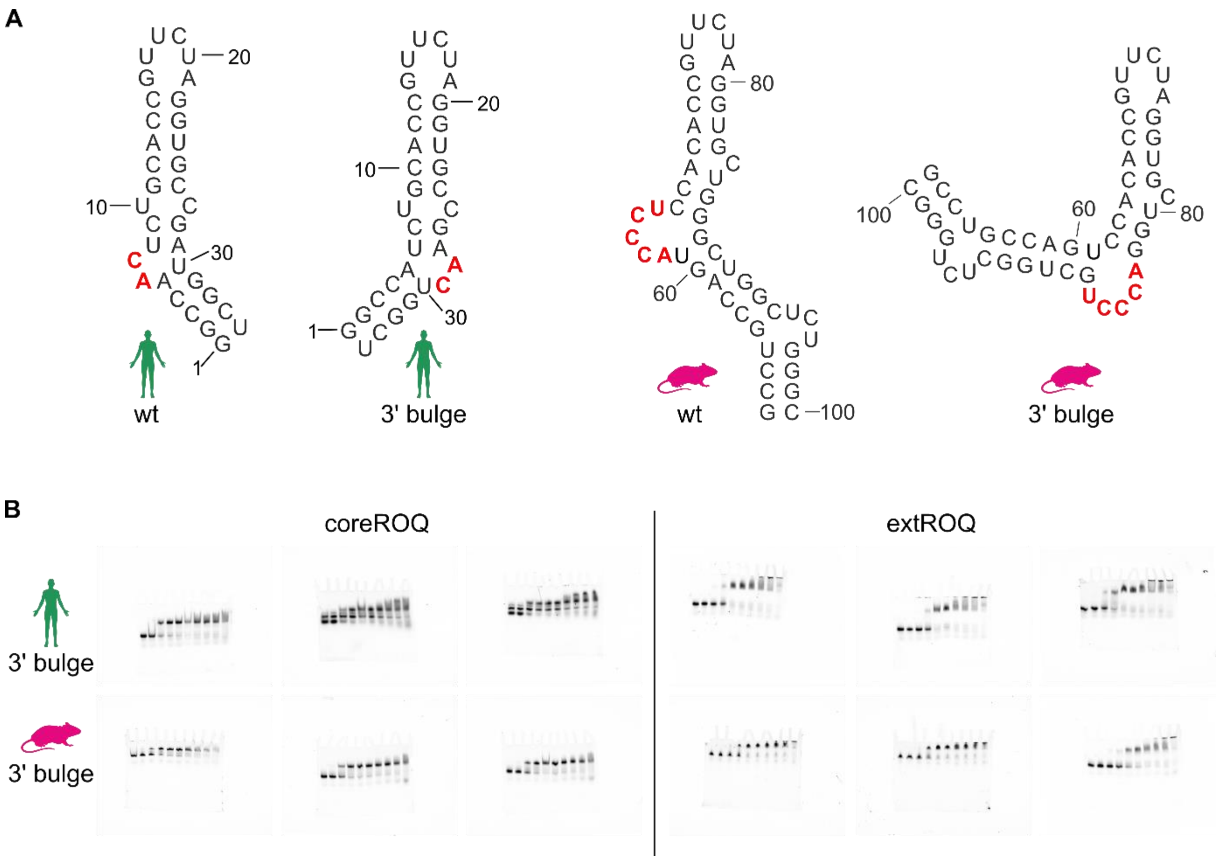


**Supplementary Figure 9.** Bulge geometry impacts complex formation with Roquin. **A)** Secondary structure predictions of human and mouse wt ADEs and 3' bulge variants. **B)** EMSAs of coreROQ (left) and extROQ (right) with human and murine 3'bulge ADE variants. Shown are triplicates. Calculated $K_D$ values are averages and plotted in Figure 6C and listed in Table 2.

**Supplementary Figure 10:**

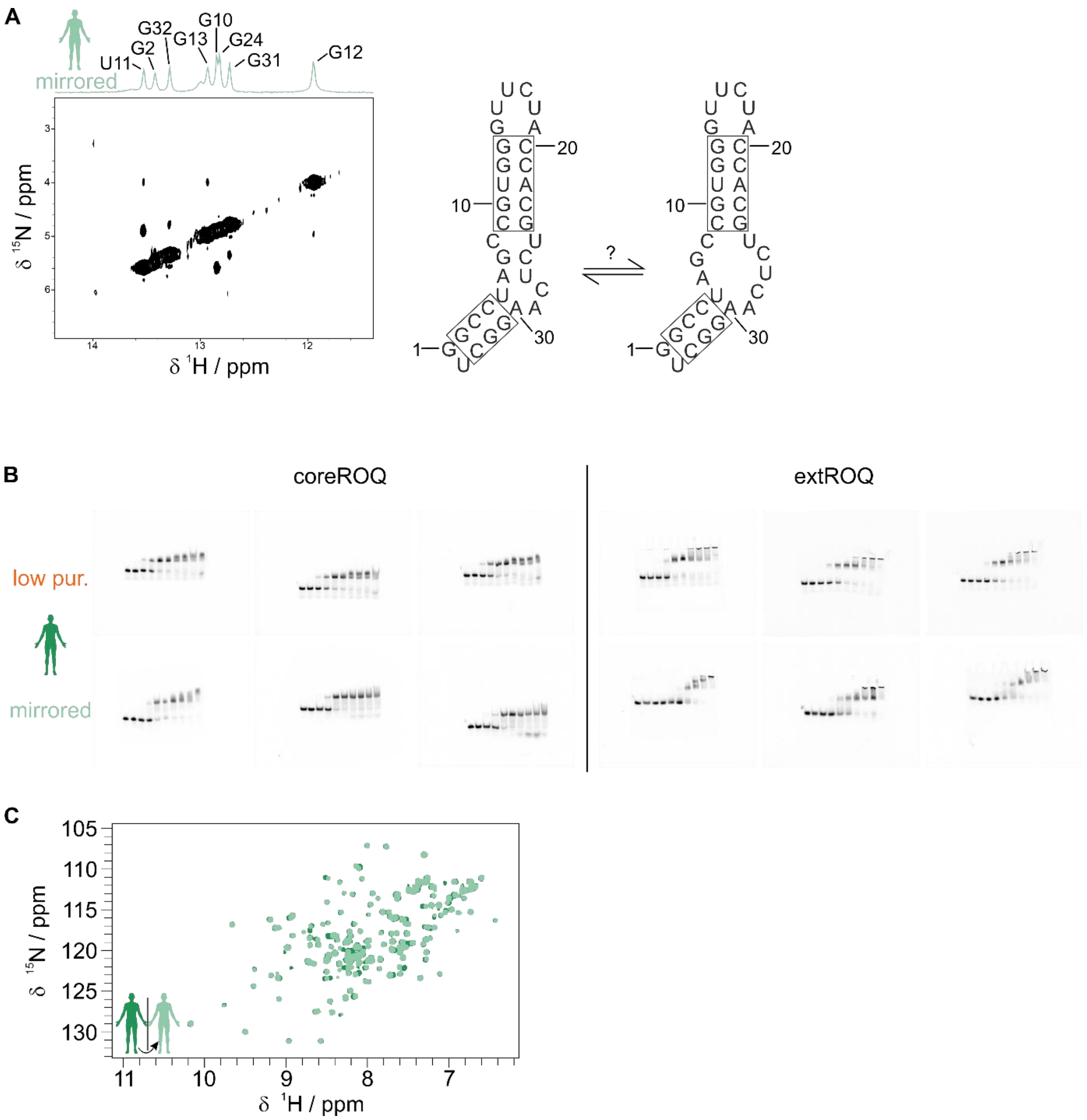


**Supplementary Figure 10.** Alterations in ADE geometry affect Roquin binding. **A)** $^{1}$H,$^{1}$H-NOESY spectrum of the mirrored human ADE. The imino proton resonance assignment is shown in the corresponding 1D spectrum on top. Confirmed base pairs are boxed in the secondary structure prediction on the right. Based on the assignment, a second conformation could form, with a partially opened bulge. **B)** EMSAs of coreROQ (left) and extROQ (right) with human ADE variants. Shown are triplicates. Calculated $K_D$ values are averages and plotted in Figure 7C and listed in Table 2. **C)** Overlay of $^{1}$H,$^{15}$N-HSQC spectra of coreROQ in complex with human wt and mirrored ADEs (dark and light green).
